# Sodium tungstate promotes vascularization to support beta cell replacement in diabetes

**DOI:** 10.64898/2026.03.09.710481

**Authors:** Ainhoa García-Alamán, Marta Fontcuberta-PiSunyer, Jonna M Saarimäki-Vire, Nea Asumaa, Marta Perea-Atienzar, Rebeca Fernandez-Ruiz, Hugo Alves-Figueiredo, Christophe Broca, Joan-Marc Servitja, Ramon Gomis, Diego Balboa, Josep Vidal, Rosa Gasa

## Abstract

Insufficient vascularization remains a major obstacle to the success of cell-based therapies for diabetes. Building on prior findings that loss of the phosphatase PTP1B enhances VEGFA production and improves graft vascularization, we investigated sodium tungstate (NaW), a pharmacological phosphatase inhibitor, as a strategy to improve transplantation outcomes. Using human fibroblast-derived insulin-producing cells and human stem cell-derived islets transplanted into the anterior chamber of the eye in immunodeficient mice, we show that NaW treatment significantly increases both vascularized area and insulin-positive tissue area, while reducing apoptosis within transplanted cells. Mechanistically, NaW upregulates VEGFA expression in transplanted cells and amplifies VEGFA-induced endothelial cell proliferation, migration, and tubulogenesis via MAPK/ERK signalling. These dual effects, which encompass stimulating both endocrine and endothelial compartments, lead to enhanced integration and function of transplanted cells. Importantly, the pro-angiogenic effects of NaW occur independently of exogenous endothelial cell supplementation, relying solely on the host’s endogenous endothelial cells. These findings position NaW, and potentially other phosphatase inhibitors, as promising adjuncts to improve vascularization, survival, and therapeutic efficacy in clinical cell-based transplantation protocols for diabetes.

**One-sentence summary:** Proangiogenic role of sodium tungstate in transplantation

## Introduction

The replacement of islet beta cells, whether sourced from donors or generated *in vitro* as islet/beta-like cells, presents promising prospects for curing type 1 diabetes (T1D) and other forms of severe insulin-dependent diabetes. Nonetheless, achieving the full therapeutic potential of these approaches requires improvements in transplantation methodologies. One major challenge is delayed and insufficient graft vascularization which results in poor cell survival and function post-transplantation. The success of engraftment heavily depends on overcoming this challenge.

Vascularization is essential for islet survival and function as it enables rapid sensing of blood glucose and coordinated release of insulin and glucagon into the circulation(*1*). For donor islets, the disruption of vascular connections during isolation, combined with the gradual depletion of intra-islet endothelial cells (iECs) during pre-transplantation islet culture (*2, 3*), is a major challenge. Following transplantation, islets must re-build a functional vascular connection with the host, a process that typically takes two to four weeks(*4*). This delay in vascularization leads to a period of oxygen and nutrient deprivation for the newly transplanted islets, both of which can affect cell function and viability and lead to cell damage and/or death. Islets are specially sensitive to hypoxic conditions, with approximately 50%–70% of transplanted cells being lost during the isolation, culture, and peri-transplantation stages of treatment(*5*). This early loss of islet mass ultimately jeopardizes the long-term success of the graft.

On the other hand, the shortage of donor islets for transplantation has prompted the development of *in vitro* strategies to generate insulin-producing cells, including stem cell–derived islets (SC-islets) from embryonic or induced pluripotent stem cells, as well as through direct reprogramming of somatic cells. These laboratory-generated islet/beta-like cells have no vasculature which poses an even greater challenge for achieving effective graft vascularization in such therapies. Encapsulation devices, designed to shield transplanted cells from immune rejection and eliminate the need for immunosuppressive therapy, often block vascular integration, exacerbating ischemia and contributing to poor graft outcomes, factors that have caused some clinical trials to be discontinued(*6*). Additionally, trials using semipermeable devices have reported inadequate and uneven engraftment of transplanted cells within the subcutaneous space, an environment characterized by low vascular density(*7–9*). To overcome these limitations, strategies such as co-transplantation with endothelial cells (ECs) and other supportive cell types(*10, 11*), or the use of microvascular fragments derived from adipose tissue(*12, 13*), have been investigated with encouraging results. Importantly, enhanced vascularization not only improves graft survival but also aids in the *in vivo* maturation of lab-generated islets(*14*). Nonetheless, the development of stable, safe, and functional autologous vascular tissue remains a critical hurdle that must be overcome to enable successful clinical translation.

The establishment of new, functional vasculature within the graft requires the activation and recruitment of ECs from both the transplant recipient and iECs within the transplanted islets (*15, 16*). The primary pro-angiogenic signal secreted by islet cells, including beta and alpha cells, is vascular endothelial growth factor A (VEGFA)(*17*). As a result, VEGFA emerged as a key candidate factor for enhancing vascularization and improving islet transplantation outcomes. Nonetheless, optimizing vascularization through external VEGFA delivery remains challenging, as precise dosing is critical: excessive angiogenesis can impair islet architecture and contribute to beta cell loss(*18–20*). In a previous study, we demonstrated that islets from mice lacking protein tyrosine phosphatase 1B (PTP1B) showed improved graft revascularization, better survival, and enhanced function, leading to improved transplant outcomes(*3*). We attributed this effect to the increased production and secretion of VEGFA by the transplanted PTP1B-deficient beta cells(*3*), highlighting the possibility of boosting endogenous pro-angiogenic pathways in cell implants as a means to accelerate their engraftment.

Building on this evidence, here we sought to determine whether a pharmacological approach could replicate these effects, providing a non-genetic more clinically translatable alternative. For this purpose, we selected tungstate, an oxyanion of tungsten known to function as a broad phosphatase inhibitor. Notably, tungstate effectively inhibits PTP1B with an IC_₅₀_ of 210 µM(*21, 22*). Among its various formulations, sodium tungstate (Na_2_WO_4_, termed NaW henceforth) stands out due to its oral bioavailability, favorable pharmacokinetic properties, and a well-established safety profile. NaW has been broadly studied for its anti-diabetic and anti-obesity effects in both preclinical models and clinical trials, with no reported toxicity or adverse effects(*23–29*).

In this study, we evaluated the potential of NaW to improve graft vascularization in the context of *in vitro* generated cell-based therapies for diabetes. For these experiments, we utilized iBeta-like cell spheroids generated via direct transcription factor-mediated reprogramming of human fibroblasts(*30*), as well as a clinically advanced beta cell replacement platform employing a well-established model of stem cell–derived islets (SC-islets), produced through stepwise differentiation of human pluripotent stem cells(*31*). Our results show that NaW significantly enhances early functional vascularization in both models, resulting in improved graft integration and overall transplantation outcomes. Importantly, this pro-vascular effect appears to be mediated through dual actions, targeting both the transplanted endocrine cells and the host endothelial cells, thereby supporting vascular remodeling and successful engraftment.

## RESULTS

### Inclusion of HUVEC cells improves vascularization of human iBeta-like cell transplants

To better recapitulate the microenvironment of donor human islets, human umbilical vein endothelial cells (HUVECs) were incorporated into iBeta-like cells during the aggregation phase of the reprogramming protocol at a 1:9 ratio (HUVEC:iBeta-like cells) (Figure 1A). After six days of culture, the resulting HUVEC-containing iBeta-like spheroids (hereafter referred to as iBeta-like/HUVEC) exhibited viability and gross morphology comparable to spheroids composed exclusively of iBeta-like cells (Figure S1A,B).

**Figure 1.**
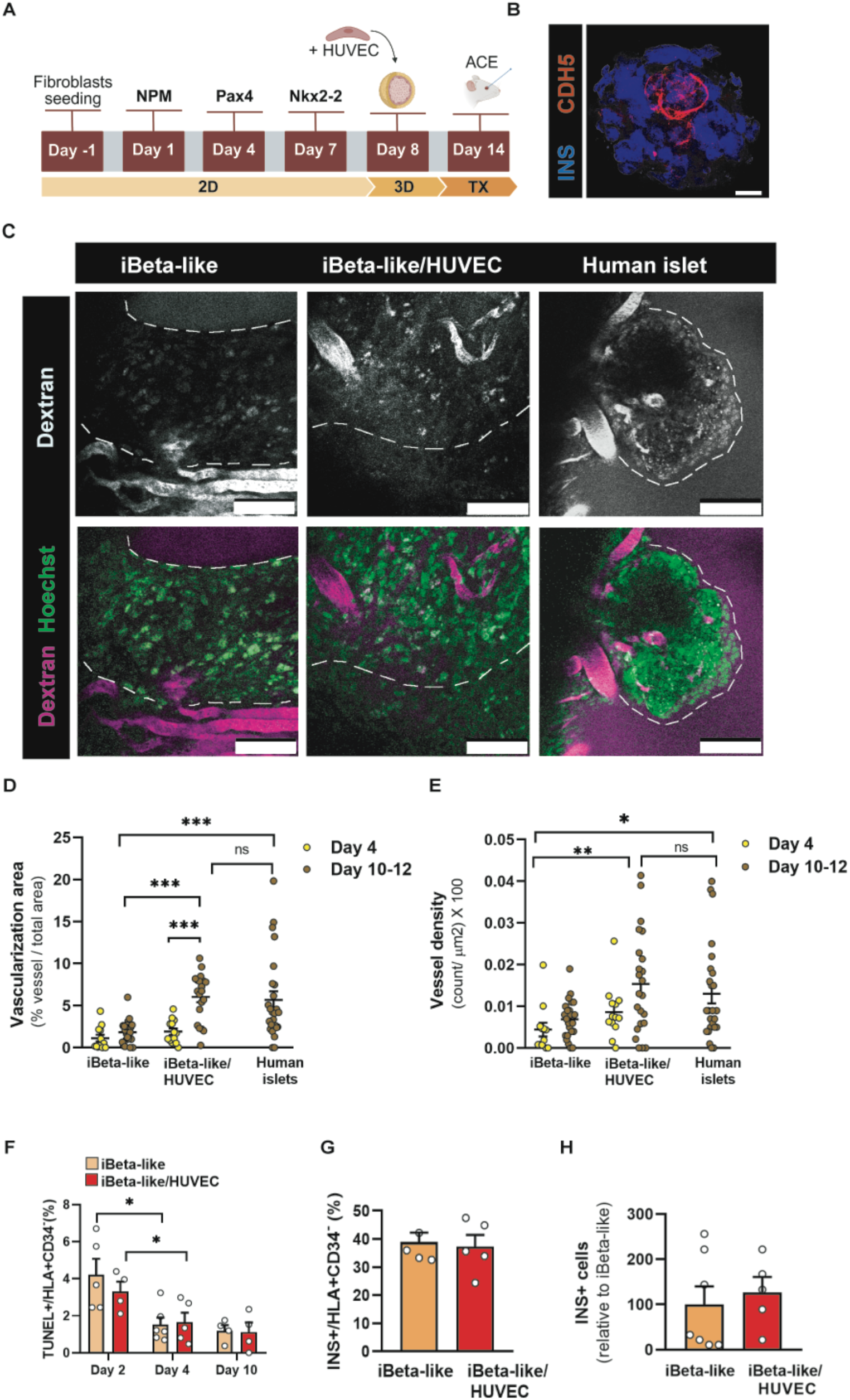
Vascularization and histological characterization of iBeta-like and iBeta-like/HUVEC grafts. **(A)** Schematic diagram of the direct reprogramming protocol used to generate iBeta-like/HUVEC cells spheroids. At day 14, spheroids (with or without HUVEC) were transplanted into the ACE of NSG mice. **(B)** Representative confocal image of an iBeta-like/HUVEC spheroid stained for insulin (INS, blue) and VE-cadherin (CDH5, red). Scale bar: 25 μm. **(C)** Representative *in vivo* two-photon microscopy images illustrating functional vasculature labelled with RITC-dextran and transplanted cells visualized via Hoechst-stained nuclei in iBeta-like and iBeta-like/HUVEC grafts at day 10 post-transplantation, and in human islet grafts at day 12 post-transplantation. Discontinued lines represent graft borders. Scale bars: 100 μm. **(D,E)** Quantitative analysis of vascularized area (D) and vessel density (E) in the indicated graft types and times after transplantation. **(F)** Quantification of cell death using TUNEL assay at the indicated time points. Data are presented as the percentage of TUNEL+ cells relative to the total transplanted cell population, excluding HUVECs. **(G)** Reprogramming rates at day 10 post-transplantation, expressed as the percentage of INS+ cells relative to the total number of human (HLA+) non-endothelial (CD34-) cells. **(H)** Quantification of INS+ cells in the indicated graft types at day 30 post-transplantation. Data are expressed relative to iBeta-like cells, with values set to 100%. Data are shown as mean ± SEM, with the number of observations indicated by individual dots. In panels D and E, each dot represents one measured area, with data collected from n = 4–6 grafts from each cell type. In panels F, G, and H, each dot represents the mean value for a single graft. Statistical significance was assessed using two-way ANOVA in panels D, E, and F for comparisons between iBeta-like and iBeta-like/HUVEC grafts across the different days analyzed; one-way ANOVA for comparisons between graft types at days 10–12 (panels D and E); and unpaired Student’s *t*-tests for panels G and H: : *p < 0.05, **p < 0.005, *** p<0.0001 for comparisons between indicated conditions.

Immunofluorescence analysis of iBeta-like/HUVEC spheroids revealed a distinct spatial organization, with HUVECs preferentially localizing to the spheroid core in a ring-like arrangement, while insulin-positive (INS^⁺^) cells were predominantly found at the periphery (Figure 1B). Gene expression analysis demonstrated comparable levels of beta cell and fibroblast-associated markers between iBeta-like and iBeta-like/HUVEC spheroids, with the exception of a significant upregulation of the prohormone convertase gene *PCSK1* in iBeta-like/HUVEC spheroids (Figure S1C). Notably, *PCSK1* mRNA was readily detected in HUVECs, which may account for the observed increase in *PCSK1* expression in iBeta-like/HUVEC spheroids (Figure S1C). Quantification of INS⁺ cells revealed no significant differences between both spheroid types (Figure S1D). Moreover, functional assessment by glucose-stimulated insulin secretion showed comparable responses between the two groups (Figure S1E). Collectively, these data indicate that the inclusion of 10% HUVECs during the 3D reprogramming stage did not markedly affect the efficiency of *in vitro* fibroblast conversion into iBeta-like cells.

We next evaluated the extent of vascularization in iBeta-like and iBeta-like/HUVEC grafts in comparison with isolated human donor islets. Transplantations were performed into the anterior chamber of the eye (ACE) of normoglycemic NSG mice, a site that enables longitudinal *in vivo* imaging and direct assessment of functional vascular integration. Vascularization was analyzed at 4 and 10 days post-transplantation for iBeta-like grafts and at day 12 for human islets using RITC-dextran, a fluorescent blood tracer that allows real-time visualization of perfused vessels within the grafts. Although no significant differences were detected at day 4, by day 10 grafts generated from iBetalike/HUVEC spheroids displayed a significantly higher fractional vascularization and vessel density compared with grafts composed of iBeta--like spheroids alone (Figure 1C-E). Notably, the extent of vascularization approached that observed in transplanted human donor islets (Figure 1C-E).These findings demonstrate that the inclusion of HUVECs enhances *in vivo* vascularization of iBeta-like grafts.

To evaluate the impact of enhanced vascularization on post-transplantation cell survival, TUNEL staining was conducted on days 2, 4, and 10. Both iBeta-like and iBeta-like/HUVEC spheroids showed a similar decrease in cell death from day 2 to day 4, which remained stable through day 10 (Figure 1F). These results suggest that the presence of HUVECs does not markedly affect graft survival during the early post-transplantation period, nor at later stages when differences in vascularization become significant. Histological analysis of the grafts revealed similar percentage of INS+ cells among HLA+CD34-(=human, non-endothelial) cells revealing no changes in reprogramming rates between the two groups (Figure 1G). Additionally, the proportion of proliferating INS^⁺^ cells remained low and varied across grafts, but did not differ significantly between groups at either day 4 or day 10 post-transplantation (Figure S2). Consistent with prior findings in control grafts(*30*), long-term evaluation up to 30 days post-engraftment showed a low number of remaining INS+ cells, with no significant differences observed between both types of grafts (Figure 1H).

Overall, these results demonstrate that the addition of HUVECs positively impacts graft vascularization, as evidenced by enhanced vascular network compared to controls. Despite this improved vascularization, there is no evident impact on the persistence or expansion of reprogrammed cells over time.

### Effects of NaW on human iBeta-like/HUVEC grafts

Given that the addition of endothelial cells alone did not markedly improve graft survival, we investigated whether combining this approach with the PTP1B inhibitor NaW could enhance outcomes, based on our previous findings showing beneficial effects of PTP1B deletion in transplanted islets(*3*). To test this hypothesis, we conducted a new series of transplantation experiments using iBeta-like/HUVEC spheroids. Post-transplantation, recipient animals were divided into two groups: one received no treatment (control), while the other was given NaW at a concentration of 1 g/L in their drinking water. Consistent with previous studies, no significant differences in body weight or glycemia were observed over the 30-day follow-up period (Figure S3).

Functional vascularization was evaluated on day 10 post-transplantation. In NaW-treated mice, the vascularization area was increased two-fold compared to untreated mice (Figure 2A,B), indicating a larger area occupied by vessels. Additionally, vessel density was elevated by 50% in the NaW group (Figure 2A,C), representing a richer microvascular network within the graft. Histological analysis revealed the presence of both host-derived mouse ECs and transplanted HUVEC cells within the grafts of NaW-treated animals. In contrast, in control animals, mouse ECs were found only in the vicinity of the grafts but not within them (Figure S4). These observations suggest that NaW enhances the recruitment of recipient ECs to the transplantation site.

**Figure 2.**
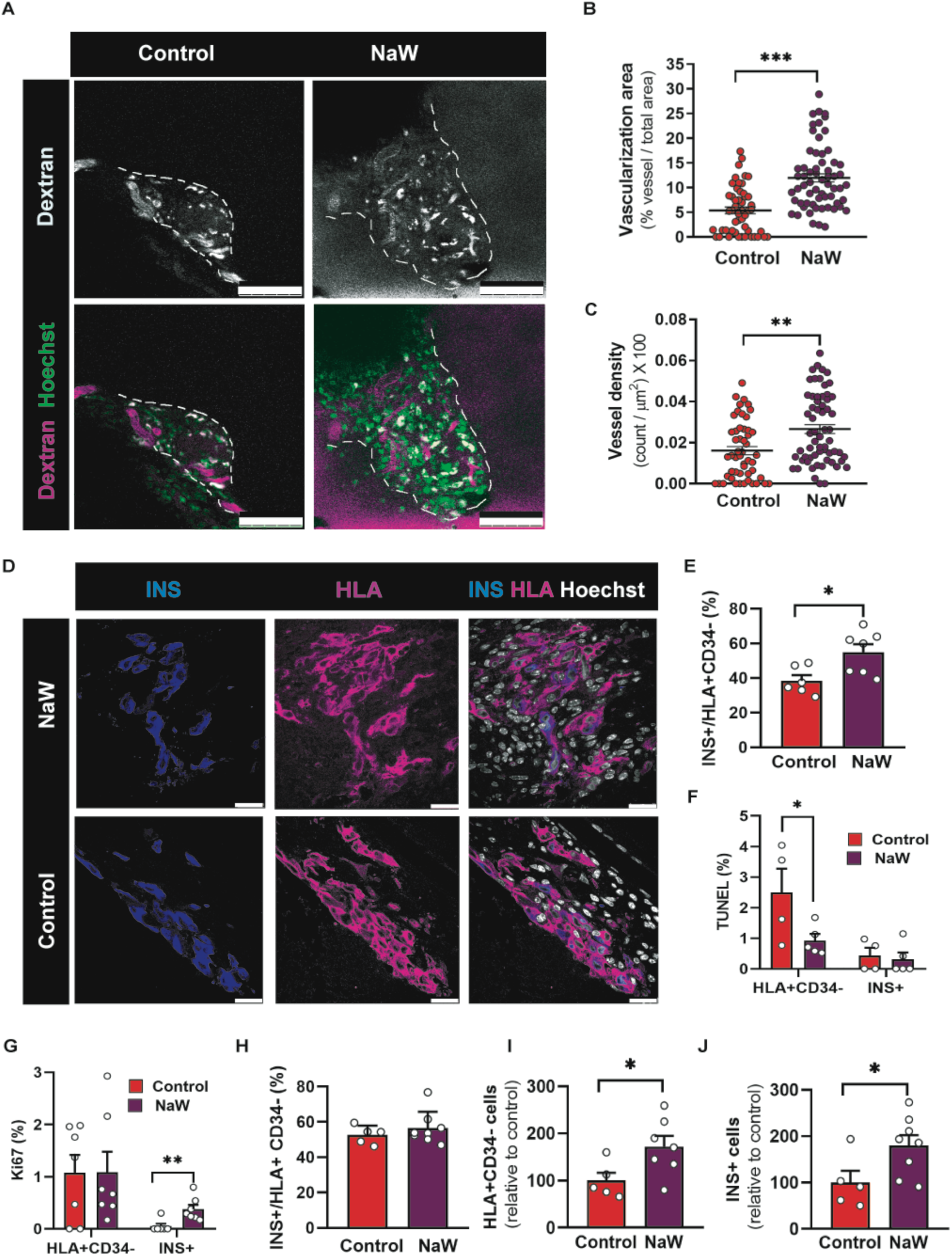
Effects of NaW treatment on iBeta-like/HUVEC grafts. **(A)** Representative *in vivo* two-photon microscopy images of iBeta-like/HUVEC grafts illustrating functional vasculature labelled with RITC-dextran and transplanted cells visualized via Hoechst-stained nuclei in control and NaW-treated mice at day 10 post-transplantation. White fluorescence observed in some cells, unrelated to vascular structures, corresponds to residual Cherry+ expression from the reprogramming adenovirus. Discontinued lines represent graft borders. Scale bars: 100 μm **(B,C)** Quantitative analysis of vascularized area (B) and vessel density (C) in grafts at day 10 post-transplantation. **(D)** Representative immunofluorescence images of grafts stained for insulin (INS), HLA (to mark human cells) and Hoechst (nuclei) in control and NaW-treated mice at day 10. Scale bar: 25 μm. **(E)** Reprogramming rates at day 10 post-transplantation, expressed as the percentage of INS+ cells relative to the total number of human (HLA+) non-endothelial (CD34-) cells. **(F,G)** Quantification of cell death by TUNEL (F) and cell proliferation by Ki67 (G) immunostaining in grafts at day 10. Percentages are shown for HLA+CD34-cells and for INS+ cells, relative to total HLA+CD34- and total INS+ cells, respectively. **(H)** Reprogramming rates at day 30 post-transplantation, expressed as the percentage of INS+ cells relative to the total number of human (HLA+) non-endothelial (CD34-) cells. **(I,J)** Quantification of HLA+CD34-cells (I) and INS+ cells (J) in grafts from control and NaW-treated mice at day 30 post-transplantation. Values are expressed relative to control grafts, with values set to 100%. Data are presented as mean ± SEM, with the number of observations indicated by individual dots. Each dot represents the mean value from a single graft, except in panels B and C, where each dot corresponds to one measured area from a total of 10–12 grafts. Statistical significance was determined using an unpaired Student’s t-test: *p < 0.05, **p < 0.005, *** p<0.0001 for comparisons between indicated conditions.

We next examined how NaW treatment influenced iBetalike cell graft outcomes. NaWtreated animals exhibited nearly a 50% increase in the proportion of INS⁺ cells among total HLA^⁺^CD34^⁻^ cells (Figure 2D,E). TUNEL assays showed a significant reduction in cell death within the HLA^⁺^CD34^⁻^ compartment in NaWtreated grafts compared to controls (Figure 2F). The percentage of INS^⁺^TUNEL^⁺^ cells was low in both groups, suggesting that the survival benefit conferred by NaW primarily affects non or partially reprogrammed cells. Overall proliferation rates within the HLA^⁺^CD34^⁻^ population were low (∼1%) and were not altered by NaW treatment. In contrast, INS^⁺^ cells in control grafts displayed almost no proliferation, whereas NaWtreated mice showed a significant, although still low (<0.5%), increase in INS^⁺^ cell proliferation (Figure 2G). Together, these findings indicate that the higher proportion of INS^⁺^ cells observed at day 10 in NaW-treated mice is more likely due to increased proliferation of reprogrammed cells rather than reduced cell death.

Finally, to evaluate the longer-term impact of NaW on graft outcomes, selected transplants were analyzed up to 30 days post-transplantation. Grafts from NaWtreated animals displayed reprogramming rates comparable to controls at day 30 (Figure 2H), remaining close to the ∼60% level already observed at day 10. Notably, while the reprogramming rate in untreated animals increased between days 10 and 30, -NaWtreated grafts had already reached a higher level at day 10 and maintained it thereafter, indicating that the effects of NaW occur early after transplantation. Moreover, by day 30, NaW-treated grafts exhibited-a two-fold increase in HLA^⁺^CD34^⁺^ and INS^⁺^ cell numbers compared with controls (Figure 2I,J), consistent with improved longterm survival of transplanted cells. Together, these findings highlight NaW as an effective -earlyacting-enhancer of graft preservation and stability.

### Effects of NaW on VEGFA production by human iBeta-like cells

In our previous study(*3*), we demonstrated that transplanted beta cells deficient in the phosphatase PTP1B produce higher levels of VEGFA, which was proposed as the mechanism responsible for enhanced graft vascularization. To determine whether the pro-vascularization effect induced by NaW in iBeta-like cell grafts was mediated by upregulation of VEGFA, we performed immunofluorescence analysis on grafts collected 10 days post-transplantation. Quantitative analysis showed that 69% of INS^⁺^ cells in NaW-treated grafts co-expressed VEGFA, compared with 43% in control grafts (Figure 3A,B). Increased VEGFA expression was also observed in insulin-negative cells, with 55% of VEGFA^⁺^/INS^⁻^ cells in NaW-treated grafts versus 23% in control grafts.

**Figure 3.**
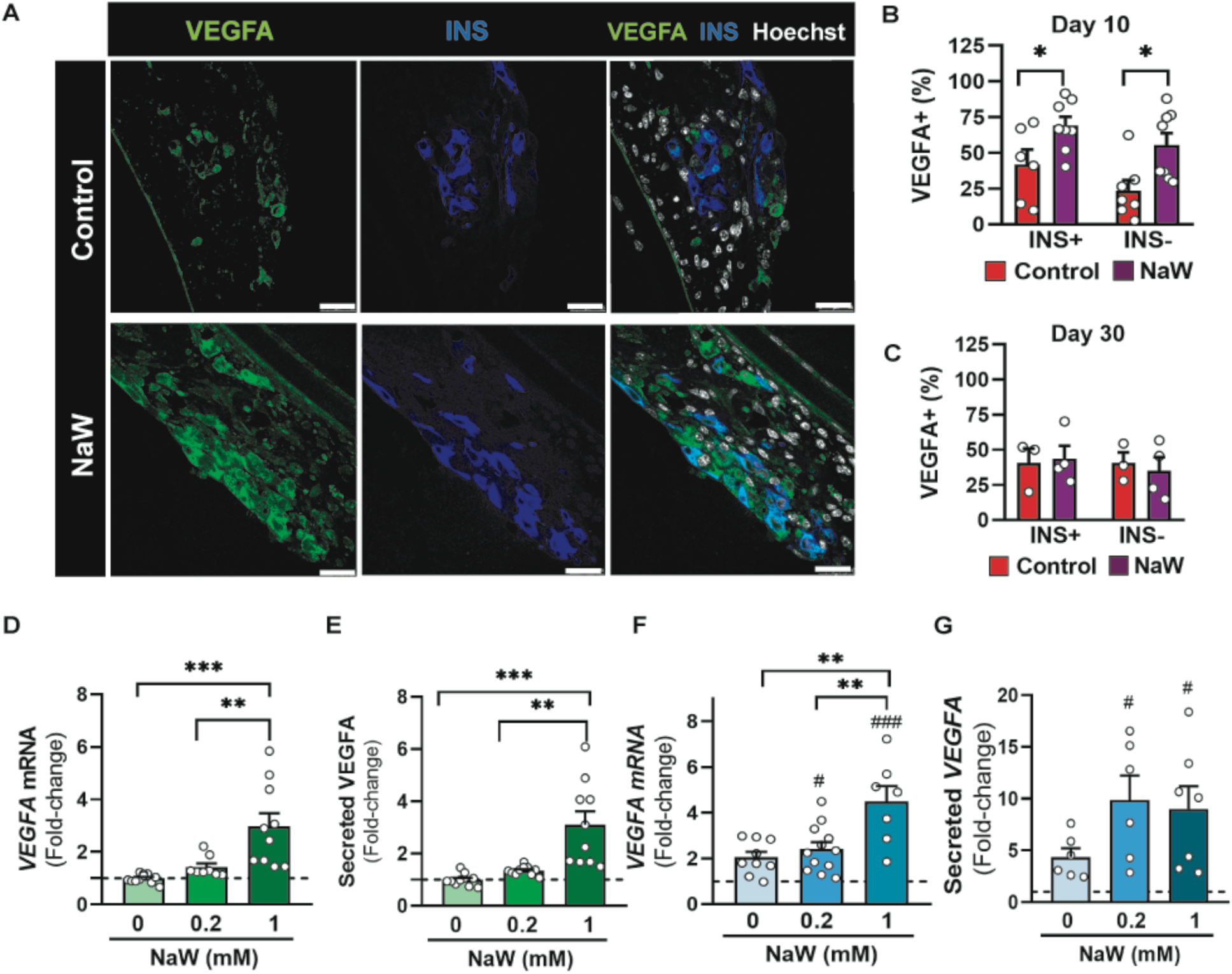
Analysis of VEGFA production by iBeta-like cells *in vivo* and *in vitro*. **(A)** Representative confocal images of iBeta-like/HUVEC grafts at day 10 post-transplantation from control and NaW-treated mice. Grafts were stained for VEGFA (green), insulin (INS, blue), and Hoechst (white, nuclei). Scale bar: 25 μm. **(B,C)** Quantification of the percentage of INS- and INS+ cells co-expressing VEGFA relative to total HLA+CD34-cells in iBeta-like/HUVEC grafts at day 10 (B) and day 30 (C) post-transplantation from control and NaW-treated mice. **(D,E)** *VEGFA* mRNA expression (D) and VEGFA protein released to the culture media (E) after culturing iBeta-like cells with or without NaW at the indicated concentrations for 48 hours. Expression was measured by qPCR using *TBP* as a housekeeping gene and the amount of secreted VEGFA protein was quantified by ELISA. Data are expressed as fold-change relative to untreated cells. **(F,G)** *VEGFA* mRNA expression (F) and VEGFA protein released to the culture media (G) after culturing iBeta-like cells with or without NaW at the indicated concentrations for 24 hours under hypoxic conditions. Expression was measured by qPCR using *TBP* as a housekeeping gene and the amount of secreted VEGFA protein was quantified by ELISA. Data are expressed as fold-change relative to untreated normoxic control. Data are presented as mean ± SEM, with individual dots indicating the number of observations. In panels B and C, each dot corresponds to a single graft. For all other panels, values are derived from at least three independent reprogramming experiments. Statistical significance was assessed using unpaired Student’s t-test between indicated groups (B,C), or one-way ANOVA (D-G): *p < 0.05, **p < 0.005, **\*\*\***p < 0.0001 for comparisons between indicated conditions; #p < 0.05 and ###p < 0.0001 relative to normoxia.

By day 30 post-transplantation, the proportions of both VEGFA^⁺^/INS^⁺^ and VEGFA^⁺^/INS^⁻^ cells were comparable between treatment groups (Figure 3C), indicating that the effect of NaW on VEGFA expression is transient. Together, these results suggest that NaW temporarily enhances VEGFA production in transplanted cells, a response that may be critical for promoting early graft vascularization.

Next, to determine whether NaW directly influences VEGFA production in iBeta-like cells, we cultured the cells with or without two different NaW concentrations. After 48 hours, we assessed *VEGFA* mRNA levels and found a significant increase in its expression in cells exposed to the highest NaW concentration (Figure 3D). Consistently, we also observed elevated levels of VEGFA protein released into the culture medium (Figure 3E). These findings demonstrate that NaW can directly enhance VEGFA production in ibeta-like cells. Next, to mimic the transplantation microenvironment, we repeated the experiment under hypoxic conditions (3% O_2_). After 48 hours, *VEGFA* mRNA and secreted protein levels were higher under hypoxia than normoxia, but the differences were not significant. In the presence of NaW, these differences became statistically significant. Moreover, 1 mM NaW markedly increased *VEGFA* mRNA expression compared to hypoxia alone (Figure 3F). VEGFA protein secretion also tended to rise with NaW, but these changes were not statistically significant when compared to hypoxia alone (Figure 3G).

Together, these findings support a direct influence of NaW on VEGFA production by iBeta-like cells, identifying VEGFA as a likely target and mediator of NaW-induced vascularization.

### Direct effects of NaW on human endothelial cells

VEGFA signalling via VEGFR2 is negatively regulated by PTP1B, which dephosphorylates VEGFR2 and thereby attenuates downstream signalling (*32*). Given this regulatory relationship, we next investigated whether NaW, in addition to stimulating VEGFA production by iBeta-like cells, could potentiate the paracrine effects of VEGFA on endothelial cells. To this end, HUVECs were treated with NaW in the presence or absence of exogenous VEGFA, and endothelial responses were assessed using standard functional assays.

Cell viability, measured by MTT assay as an indicator of mitochondrial metabolic activity, was comparable across all treatment groups (Figure 4A), confirming that NaW was not cytotoxic under the conditions tested. In contrast, direct cell counts, which quantify actual increases in cell number, revealed a significant rise in proliferation in NaWtreated HUVECs, but only in the presence of VEGFA (Figure 4B). These data indicate that NaW enhances VEGFAdriven proliferative signalling.

**Figure 4.**
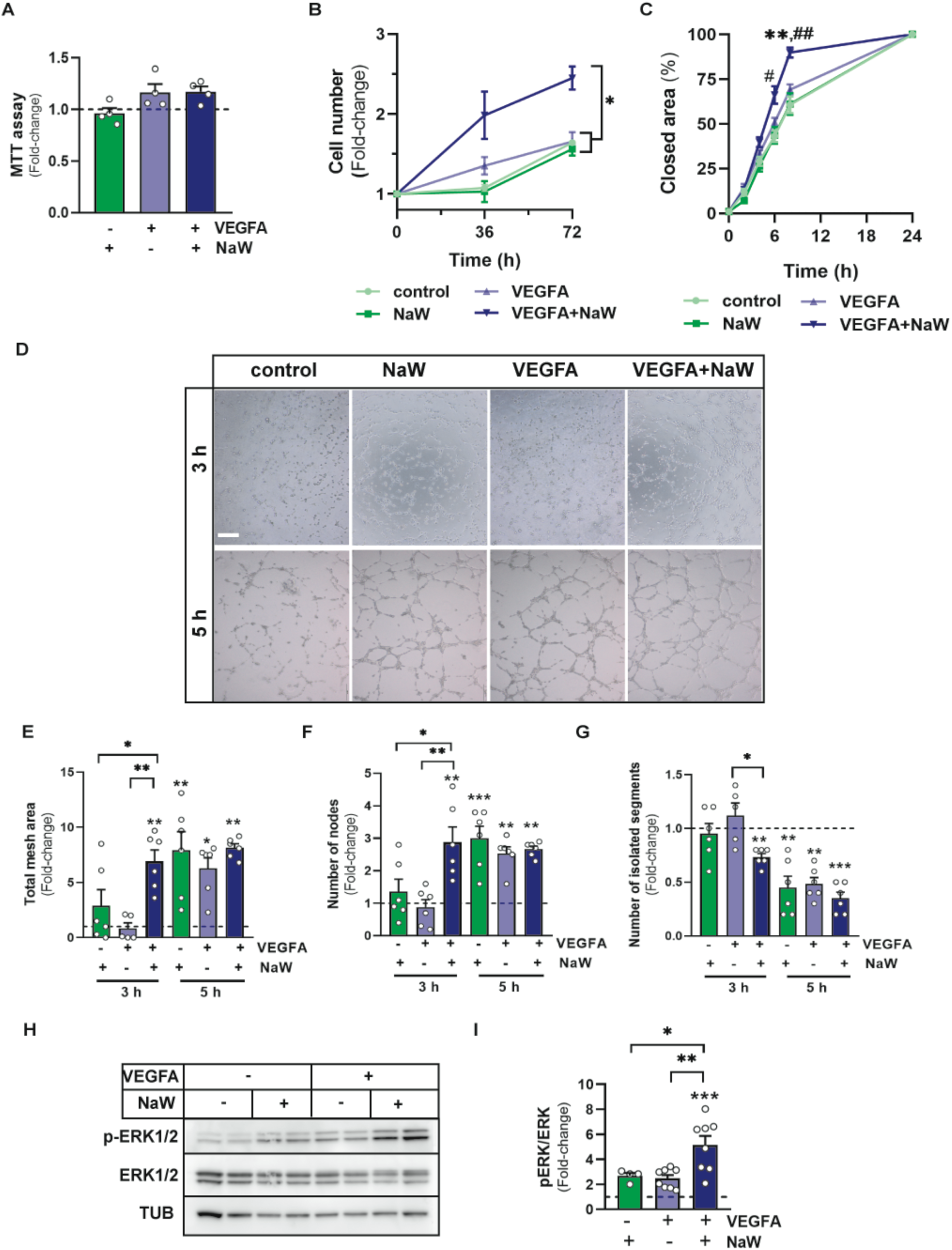
Effects of NaW on endothelial cell function *in vitro*. **(A)** HUVEC viability assessed by MTT assay after 48 hours of culture in the indicated conditions. Results are normalized to control (no VEGFA, no NaW), set to a value of 1 (dotted line). **(B)** HUVEC proliferation evaluated by direct cell counting at the indicated time points. Data are expressed as fold-increase relative to cell number at time 0 (set to 1) for each condition. **(C)** HUVEC migration assessed by wound healing assay. Results are expressed relative to the 24-hour time point, when wound closure reached 100% in all conditions. **(D)** Representative images showing tube formation by HUVECs cultured for 3 and 5 hours under the indicated conditions. Scale bar: 200 μm. **(E-G)** Quantitative analysis of tubulogenesis at 3 and 5 hours: total mesh area (E), number of nodes (F), and number of isolated segments (G). Data are expressed as fold-increase relative to control cells (no VEGFA, no NaW), which are normalized to 1. **(H,I)** Immunoblot analysis of ERK1/2 phosphorylation following 10 minutes of stimulation with VEGFA and NaW. Tubulin (TUB) serves as loading control. Panel H shows representative blots; panel I shows quantification of band intensities. Data are normalized to control cells (no VEGFA, no NaW), with values set to 1 (dotted line). Data are presented as mean ± SEM for the number of n indicated by dots. In panel B, n=3-4 and in panel C, n=9. Statistical significances: *p < 0.05, **p < 0.005, ***p < 0.0001 between indicated conditions in panels B,E,F G,I. In panel C, **p < 0.005 relative to VEGFA; #p<0.05, ##p < 0.005 relative to control cells (no VEGFA, no NaW). Tests used are one-way (A,E,F,G,I) or two-way ANOVA (B,C).

Endothelial motility was then evaluated using a woundhealing assay, which monitors the capacity of cells to migrate and repopulate a defined gap in a monolayer. NaW significantly increased migration when VEGFA was present, whereas no effect was observed with NaW alone (Figure 4C), supporting a VEGFAdependent enhancement of motility.

To assess proangiogenic behaviour, a tubeformation assay was performed to quantify the ability of endothelial cells to assemble into capillarylike structures. A rapid and pronounced increase in tubulogenesis was detected within 3 hours in the VEGFA+NaW group (Figure 4D-G). Notably, by 5 hours, NaW alone induced tube formation to a level comparable to that achieved with VEGFA alone or with the VEGFA+NaW combination, suggesting that NaW may also independently accelerate early angiogenic organization.

To further define the signalling pathways involved, we examined activation of the MAPK/ERK1/2 cascade, a central mediator of VEGFAdriven endothelial responses. ERK1/2 phosphorylation was assessed by immunoblotting. Whereas VEGFA or NaW alone induced only modest ERK1/2 activation, cotreatment resulted in a pronounced increase in phosphorylation (Figure 4H,I), indicative of a synergistic effect. Given ERK1/2’s wellestablished role in promoting endothelial proliferation, migration, and tubulogenesis, these results suggest that NaW enhances VEGFAinduced angiogenic behaviour by potentiating MAPK/ERK signalling.

Collectively, these findings indicate that the proangiogenic actions of NaW *in vivo* arise not only from its ability to stimulate VEGFA production by iBetalike cells, but also from its capacity to amplify VEGFAdependent endothelial responses. This dual mechanism likely contributes to improved graft vascularization and may enhance the therapeutic efficacy of NaW in transplantation settings.

### Effects of NaW on human SC-islet grafts

To increase the translational relevance of these findings, we extended our experimental paradigm to human embryonic stem cell-derived islets (SC-islets), a clinically validated cell source (*33*) (Figure S5). Similar to primary human islets, SC-islet engraftment is often hindered by inadequate vascularization. By evaluating NaW in this context, we aimed to further validate its pro-vascular effects using a cell model currently employed in clinical protocols.

SC-islets were transplanted into the ACE of normoglycemic NSG mice, and *in vivo* vascularization was assessed at days 15 and 30 post-transplantation. Throughout the study, no differences in body weight or glycemia were observed between groups (Figure S6), indicating that neither the transplantation procedure nor NaW treatment affected systemic metabolic parameters.

At day 15, grafts from NaW-treated mice displayed a significantly greater vascularized area and higher vessel density compared with controls, and these differences remained robust at day 30 (Figure 5A-C). Vascular structures appeared to originate exclusively from recipient mouse endothelial cells, as CD34 immunostaining did not detect human CD34^⁺^ cells within the grafts (Figure S7). This observation is consistent with prior reports (*34*) and further supports the notion that host endothelial cells are the primary contributors to SC-islet graft vascularization under the conditions examined.

**Figure 5.**
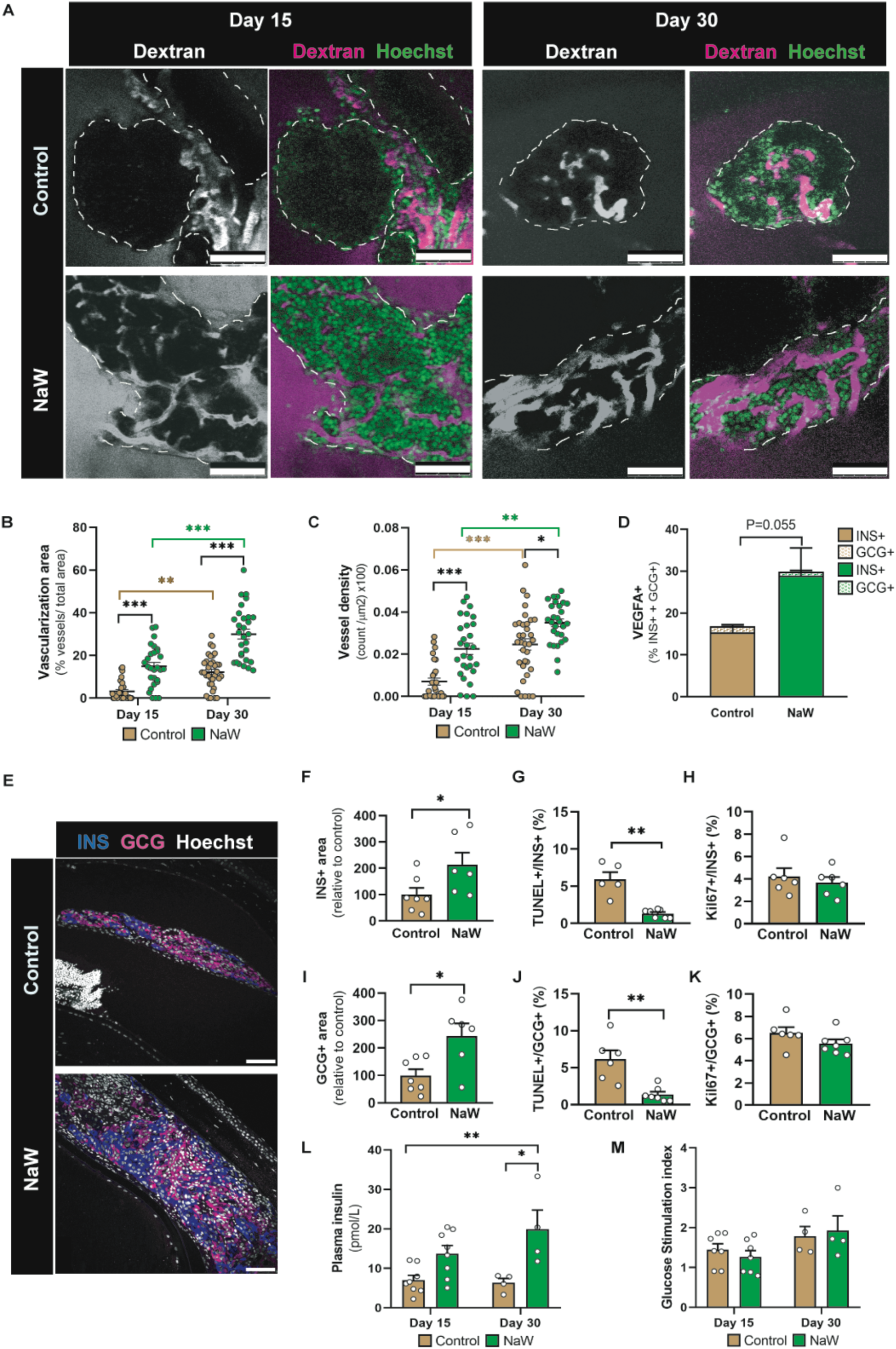
Effects of NaW treatment on SC-islet grafts. **(A)** Representative *in vivo* two-photon microscopy images of SC-islet grafts illustrating functional vasculature labelled with RITC-dextran and transplanted cells visualized via Hoechst-stained nuclei in control and NaW-treated mice at day 10 post-transplantation. Discontinued lines represent graft borders. Scale bars: 100 μm **(B,C)** Quantitative analysis of vascularized area (B) and vessel density (C) in grafts at days 15 and 30 post-transplantation. **(D)** Quantitative analysis of the percentage of endocrine cells (including INS+ and GCG+ cells) co-expressing VEGFA in grafts at day 5 post-transplantation from control and NaW-treated mice. **(E)** Representative immunofluorescence images of grafts stained for insulin (INS), glucagon (GCG) and Hoechst (nuclei) in control and NaW-treated mice at day 30. Scale bar: 75 μm. **(F,I)** Quantification of insulin-positive (F) and glucagon-positive (I) areas in grafts from control and NaW-treated mice at day 30 post-transplantation. Values are expressed relative to control grafts, with values set to 100%. **(G,J)** Quantification of cell death by TUNEL. Values represent the percentage of TUNEL+INS+(G) and TUNEL+GCG+ (J) cells relative to total INS+ and GCG+, respectively, in the indicated type of grafts at day 15 post-transplantation. **(H,K)** Quantification of cell proliferation by Ki67 staining. Values represent the percentage of Ki67+INS+ (H) and Ki67+GCG+ (K) cells relative to total INS+ and GCG+ in indicated type of grafts at day 15 post-transplantation. **(L)** Fasting human plasma insulin in control mice and mice treated with NaW at days 15 and 30 post-transplantation. **(M)** Fold-increase in plasma insulin 20 min after intraperitoneal glucose administration in control mice and mice treated with NaW at days 15 and 30 post-transplantation. Data are presented as mean ± SEM. The number of observations are indicated by individual dots, except for panel D (n=4). In panels B and C, each dot represents one measured area from a total of n = 6–9 grafts. In panels F-K, dots correspond to the mean value from individual grafts, while in panels L and M, each dot represents an individual mouse. Statistical significance was assessed using one-tailed unpaired Student’s t-test (D), two-tailed unpaired Student’s t-test (F-K), or two-way ANOVA (B, C, G, H): *p < 0.05, **p < 0.005, ***p < 0.0001 for comparisons between indicated conditions. Significance bars and symbols are color-coded for comparisons within the same treatment group across different days and shown in black for comparisons between different treatment groups.

To explore the mechanisms driving the enhanced vascularization, we performed VEGFA staining. At day 15, VEGFA protein was detected in only a few isolated cells in both treatment groups, prompting us to examine earlier time points. By day 5, both control and NaW-treated grafts exhibited a marked increase in VEGFA-positive cells. Notably, NaW-treated grafts showed a trend toward a higher proportion of VEGFA-positive endocrine cells (combined INS^⁺^ + GCG^⁺^ cell populations) (Figure 5D). Remarkably, the vast majority of VEGFA-expressing cells were insulin-positive, representing 91% of VEGFA^⁺^ cells in control grafts and 96% in NaW-treated grafts. These findings indicate that beta-like cells are the predominant source of VEGFA within SC-islet grafts at this early stage.

Quantification of the insulin-positive area at day 30 demonstrated a two-fold increase in NaW-treated mice, indicating improved graft integration (Figure 5E,F). Analysis of cell death by TUNEL staining at day 15 showed a marked reduction in INS+ cell apoptosis (Figure 5G), whereas Ki67 staining revealed no significant differences in INS+ cell proliferation (Figure 5H). Similarly, NaW-treated grafts showed an expansion of the glucagon-positive area, reduced GCG+ cell death, and unchanged GCG+ cell proliferation, suggesting that NaW effects extend beyond the beta cell compartment to support overall graft survival (Figure 5E,I-K).

Plasma human insulin levels exhibited a trend toward higher values in NaW-treated mice at day 15 and reached statistical significance by day 30 (Figure 5L). Despite these elevations in insulin, the glucose stimulation index was similar between treatment groups at both time points (Figure 5M), indicating that functional glucose responsiveness was preserved under all conditions.

Building upon our previous findings in iBetalike/HUVEC spheroids, we next evaluated the impact of NaW treatment on grafts composed of SCislets combined with exogenous endothelial cells. HUVECs were mixed with dissociated SCislet cells at a 1:9 ratio, and the resulting aggregates were allowed to form spheroids in lowattachment plates for three days prior to transplantation. NaW treatment did not affect body weight or fasting glycemia in transplanted mice (Figure S8A,B). At 15 days posttransplantation, the NaWtreated group exhibited a higher graft vessel density, but the overall graft vascularization area was not significantly different compared to untreated controls (Figure S8C,D). By day 30, NaWtreated grafts had significantly increased vascularization and vessel density relative to the untreated group (Figure S8C,D). Interestingly, at day 30, the degree of vascularization area achieved in NaWtreated SCislet/HUVEC grafts was similar to that observed in SCislet grafts treated with NaW, although vessel density was 1.6-fold higher in the presence of HUVECs. Notably, these increases in vascular area and density were not accompanied by enhancements in beta cell mass or graft function, as indicated by unchanged insulinpositive area and circulating insulin concentrations between NaWtreated and control groups at day 30 post-transplantation (Figure S8E-G). These observations suggest that beyond a certain threshold, additional vascular density does not further improve SCislet graft survival or function.

Collectively, these findings demonstrate that NaW exerts robust proangiogenic effects on SCislet grafts independent of exogenous endothelial cell addition, and that these effects likely contribute to the improved transplantation outcomes observed in NaWtreated recipient.

## DISCUSSION

Limited and delayed vascularization represents a major obstacle to the success of cell-based therapies for diabetes, as it impairs the timely delivery of oxygen and nutrients to transplanted insulin-producing cells. This deficiency compromises graft survival and function, ultimately undermining long-term therapeutic outcomes. In this study, we demonstrate that NaW, a small molecule characterized by favourable oral bioavailability and a well-established safety profile, significantly promotes early vascularization in grafts composed of either human fibroblast-derived iBeta-like cells or human stem cell-derived islets. Mechanistically, NaW exerts a dual pro-angiogenic effect: it enhances VEGFA production by the transplanted endocrine cells and simultaneously potentiates VEGF-dependent pro-vascular signalling pathways in endothelial cells. These complementary actions lead to improved graft integration and survival *in vivo,* highlighting NaW’s potential as a clinically relevant adjunct to enhance the efficacy of cell-based interventions for diabetes.

Several strategies have been proposed to accelerate vascularization, including VEGF overexpression, incorporation of accessory cell types, use of prevascularized scaffolds, and transplantation of adipose-derived microvessels. However, each approach presents significant regulatory, manufacturing, and biological challenges. VEGF overexpression demands precise control of dosage and timing to avoid excessive or disorganized angiogenesis, which can compromise graft morphology and function(*19, 20*). Cell- and scaffold-based strategies raise concerns regarding immunogenicity, scalability, and fibrotic responses. Adipose-derived microvessels(*13*), which preserve native vascular architecture and are minimally manipulated, offer promise but may be limited by reduced quality and angiogenic potential in patients with T1D(*35, 36*), as well as by complex processing and sterility requirements that hinder widespread clinical application.

By contrast, NaW is an off-the-shelf, chemically defined small molecule that is orally bioavailable, has undergone prior clinical testing for use in metabolic diseases and fertility treatments, and has demonstrated an absence of toxicity(*28, 29, 37*). This favorable profile supports rapid translation to applications where enhancing graft vascularization is required. NaW also offers practical advantages, including low cost, stability, and suitability for systemic administration, features that may facilitate clinical implementation. Moreover, its use can be readily combined with more sophisticated pro-angiogenic strategies, providing a straightforward means to further augment their efficacy.

Our data suggest that NaW promotes vascularization through two complementary pathways. First, in transplanted endocrine cells, NaW increases VEGFA production, providing a paracrine signal for endothelial ingrowth. Second, NaW directly enhances endothelial cell function, as evidenced by its effects on EC proliferation, migration, and tube formation *in vitro*. Remarkably, most of these direct effects depend on the presence of VEGFA in *in vitro* experiments, highlighting the potential of NaW as a controllable strategy to boost graft vascularization. Whether NaW influences later stages of vessel maturation, such as pericyte recruitment, basement membrane deposition, or vascular stabilization, remains to be determined. In addition, the upstream targets of NaW in ECs are not yet fully defined. While previous studies suggest PTP1B inhibition as a likely mechanism for the observed signalling amplification, other phosphatases or signaling nodes may also contribute.

Remarkably, our data show that NaW’s pro-angiogenic effects in SC-islet grafts are mediated primarily through host endothelial cells, without requiring the presence of endothelial cells in the donor islets. This finding simplifies graft preparation by eliminating the need to pre-incorporate endothelial components. Notably, treatment with NaW did not enhance insulin graft area or circulating insulin levels in mice transplanted with SC-islets containing HUVECs, despite a marked increase in vessel density compared to SC-islets without HUVECs. This apparent paradox may stem from overly robust vascularization, which can potentially reduce cell viability or disrupt the spatial arrangement of the graft (*18–20*). Additionally, adverse effects encountered during the assembly and maintenance of SC-islet + HUVEC spheroids *in vitro* may ultimately compromise both the survival and functional integration of the graft. In this study, we did not test whether adding endothelial cells at the time of transplantation, instead of during spheroid formation, might be beneficial. Further research is required to clarify these possibilities before concluding that donor endothelial cells do not provide added value beyond NaW treatment alone.

## Limitations of the study

This study primarily evaluated vascular outcomes, leveraging the ACE as a transplantation site due to its accessibility and suitability for high-resolution *in vivo* imaging. While the ACE provided clear advantages for assessing early graft vascularization, its translational relevance is limited. Although ongoing clinical trials are investigating the ACE as a potential site for donor islet transplantation in patients with type 1 diabetes (NCT04198350, NCT02846571, NCT02916680), further investigation of NaW effects in other clinically relevant sites such as the subcutaneous, omental or muscular sites is warranted. These anatomical niches can accommodate larger graft volumes but present additional challenges in achieving adequate vascularization(*38, 39*). Importantly, the restricted graft capacity within the murine ACE site in mice precluded detailed functional analysis under hyperglycemic conditions, thereby preventing evaluation of long-term metabolic outcomes in the present study. Future work should therefore extend these observations to transplantation models with higher graft loads and longer follow-up, where direct assessment of glycemic control can be achieved.

Our data support the pro-angiogenic effects of NaW in an immunodeficient mouse model at doses previously deemed safe in preclinical studies. However, due to NaW’s broad phosphatase-inhibitory activity, there may be off-target impacts on immune cell function and fibrotic remodeling that require careful evaluation. Future investigations should focus on determining the optimal timing of NaW administration, with an emphasis on transient use during the peri-transplantation period, as its effects on vascularization are expected to occur early. Alternatively, exploring localized delivery methods, such as nanoparticles, hydrogels, scaffolds, or direct co-formulation with grafts, could help concentrate NaW’s effects within the graft microenvironment while reducing systemic exposure. Additionally, repurposing more selective, clinically tested phosphatase inhibitors, several of which have progressed to early-phase clinical trials(*40, 41*), may offer an alternative translational route, potentially speeding up the clinical development of this therapeutic approach.

## Conclusions

In conclusion, our study identifies sodium tungstate as a promising pharmacological agent to enhance graft revascularization in human islet-like cell transplantation. By acting on both donor-derived endocrine cells and host endothelial cells, NaW facilitates early functional vascularization, which is a key determinant of graft survival and long-term efficacy. Its favorable safety profile supports further investigation of NaW as a translatable strategy to improve the success of beta cell replacement therapies. Together with our previous findings in PTP1B-deficient mice, these results strengthen the rationale for targeting the PTP1B axis to optimize outcomes in islet and stem cell–based transplantation. More broadly, our work highlights the potential of combining soluble chemical modulators with existing and emerging biological and biotechnological approaches to advance engraftment in cell transplantation protocols.

## Materials and Methods

### Study design

The aim of this study was to determine whether sodium tungstate (NaW), a PTP1B inhibitor, promotes vascularization of beta-like cell grafts. Two cellular models were used: human fibroblast-derived beta-like cells and embryonic stem cell-derived islets, with or without co-transplantation of human umbilical vein endothelial cells (HUVECs). Grafts were transplanted into the anterior chamber of the eye of immunodeficient NSG mice and analyzed at multiple time points up to day 30. NaW was administered in drinking water post-transplantation. Functional vascularization was assessed by RITC-dextran injection and visualized using two-photon confocal microscopy. Explanted grafts were fixed and processed for immunofluorescence to quantify cell death (TUNEL), proliferation (Ki67), insulin-positive and glucagon areas, and reprogramming rates. VEGFA expression within grafts was examined to explore angiogenic mechanisms. Graft functionality was determined in SC-islet transplants by measuring plasma human insulin levels under basal conditions and following glucose challenge. Mice were randomly allocated to experimental groups and identified by unique codes. Animals that died perioperatively or exhibited severe distress (e.g., >20% weight loss or labored breathing) were euthanized by CO_₂_ inhalation and excluded. Only mice surviving the predefined timeline were included in analyses. Sample size was estimated using GRANMO software, targeting 80% power and a two-sided significance level of 5%. Data analysis was performed in a blinded manner. All procedures were approved by the Animal Ethics Committee of the University of Barcelona and conducted in compliance with FELASA guidelines, Spanish Royal Decree 53/2013, and European Directive 2010/63/EU. *In vitro* studies assessed direct effects of NaW on beta-like cells by measuring VEGFA expression and secretion, and on endothelial cells using HUVECs to evaluate viability, proliferation, migration, wound healing, and signaling pathways via immunoblotting. Each *in vitro* experiment included a minimum of three biological replicates, with exact numbers provided in figure legends.

### Mice

NOD scid gamma (NSG™) mice were obtained from Charles River Laboratories (London, England). Animals were housed under a standard 12-hour light/dark cycle with unrestricted access to food and water. Mice, males and females, aged between 8 and 20 weeks were used as recipients for transplantation experiments.

### Generation of iBeta-like/HUVEC spheroids and culture

Human foreskin fibroblasts were reprogrammed into insulin-producing beta-like cells (iBeta-like) using five transcription factors: Neurog3, Pdx1, Mafa, Pax4, and Nkx2.2, as previously described(*30*). Human umbilical vein endothelial cells (HUVEC; Life Technologies S.A.) were routinely grown in Microvascular Endothelial Cell Growth Medium-2 (EGM-2) (Lonza, Switzerland) and antibiotics.

To generate mixed iBeta-like/EC spheroids, HUVECs were incorporated at a 1:9 ratio (HUVEC:iBeta-like) at the time of spheroid formation during the reprogramming protocol. In brief, spheroids containing a total of 1200–1600 cells per spheroid were prepared one day after Ad-Nkx2.2 infection using AggreWell-800 plates (StemCell Technologies, Saint Égrève, France). Cell aggregation and subsequent 6-day culture were performed in a hybrid medium composed of 75% RPMI-1640 supplemented with 6% (v/v) fetal bovine serum (FBS), 1% (v/v) L-glutamine (Hyclone), 25% EGM-2 and antibiotics. This medium composition was initially tested to confirm cell viability and lack of interference with the reprogramming process. This proportion was chosen to roughly approximate the estimated <5% of endothelial cell content and average 10% vascular volume of native islets (*42–44*). Higher HUVEC proportions reported in the literature (*45, 46*) were also evaluated; however, 1:4 and 1:1 HUVEC:iBeta-like ratios produced spheroids with irregular morphology that failed to maintain structural integrity and progressively dissociated over the 6-day culture period.

For hypoxia experiments, iBeta-like cells were maintained in 6-well plates during 3 days after Nkx2-2 introduction and then treated with or without NaW (0.2 or 1mM) in RPMI-1640 supplemented with 6% (v/v) FBS, 1% (v/v) L-glutamine (Hyclone) and antibiotics. Cells were incubated under hypoxic conditions (3% O_₂_), achieved by regulated nitrogen supplementation, for 48 hours. Control cells were maintained under normoxic conditions (21% O_₂_) in an standard incubator. Following incubation, cell pellets were collected for gene expression analysis, and culture media were harvested for quantification of human VEGFA levels using an ELISA kit (RayBiotech, Atlanta, GA, USA).

### Human islets

Human islets were prepared by collagenase digestion followed by density gradient purification at the Laboratory of Cell Therapy for Diabetes (Hospital Saint-Eloi, Montpellier, France), as previously described. After reception, human islets were maintained in culture for 1-3 days in RPMI-1640 with 5.5mM glucose, 10% fetal bovine serum (FBS) and antibiotics, before performing the transplantation experiments. Islets from two donors (Table S1) were used for Figure 1A. Experiments involving human islets were performed in agreement with the local ethic committee (CHU, Montpellier) and the institutional ethical committee of the French Agence de la Biomédecine (DC Nos. 2014-2473 and 2016-2716). Informed consent was obtained for donors.

### SC-islet differentiation

The use of human embryonic stem-cell lines does not require a specific ethical approval according to Finnish legislation. The H1 embryonic stem-cell line (WiCell, WA01) was differentiated to stem cell-derived islets (SC-islets) with a seven-stage protocol(*1, 2*) detailed in Barsby et al(*47*). Briefly, the human embryonic stem cells were induced to differentiate through definitive endoderm, primitive gut tube and posterior foregut-like states in monolayer to pancreatic progenitors using media changes with defined small-molecule and growth-factor compositions. Two days after pancreatic progenitor stage initiation, the cells were dissociated and reaggregated using AggreWell 400 plates (Stemcell Technologies #34425). Then, aggregates were transferred to suspension culture (6-well plates, in rotation), and further differentiated through an endocrine progenitor state to insulin and glucagon positive endocrine cells (SC-islets). The final SC-islets were matured for 3-4 weeks in the final stage of the differentiation prior to being transplanted. Quality control checkpoints were performed using flow cytometry at definitive endoderm stage (determining CXCR4+ %), pancreatic progenitor stage (PDX1+NKX6.1+ %) and functional SC-islet stage (Stage 7 week 3, PDX1+NKX6.1+%, NKX6.1+CPEP+%, and INS+GCG+%), performed as previously described(*31, 47*). SC-islet functionality was assessed using dynamic glucose stimulated insulin secretion in response to low glucose (G2.8mM), high glucose (G16.8mM), stimulated with GLP1 analog Exendin 4 (G16.8mM+Ex4) and a final membrane depolarization stimulus (G2.8mM+ KCl), as detailed previously(*31, 47*).

### Transplantation into the anterior chamber of the eye

iBeta-like spheroids, SC-islets and human islets were labelled with 10 μM CFDA SE (Vybrant® CFDA SE Cell Tracer Kit, Invitrogen) for 15 min at 37^°^C before the transplant. NSG^TM^ mice were transplanted with 250-300 iBeta-like spheroids, SC-islets or human islets in the anterior chamber of the eye (ACE) as previously described(*3*). Briefly, receptor mice were anesthetized with a mix of ketamine + xylazine (80 mg/Kg + 5 mg/Kg) intraperitoneally and received a subcutaneous injection of the analgesic buprenorphine (0,1mg/Kg). Spheroids and islets were introduced in the ACE under a stereomicroscope, using a blunt polythene cannula (Fine Bore Polythene Tubing, Smiths Medical International, UK) and a Hamilton syringe. Viscofresh (10 mg/ml sodium carmellose) was used to relieve eye discomfort when the mice recovered from the anesthetics.

### Sodium tungstate treatment

After transplantation, mice were randomly assigned to receive sodium tungstate (NaW, Ref. 223336, Sigma) or no treatment. NaW was provided ad libitum in the drinking water at 1 g/L. This concentration was chosen because it is the lowest dose consistently producing biological effects without systemic toxicity. Toxicology data show that 1 g/L is well tolerated long-term, with only mild renal changes after prolonged exposure, and 28-day studies report no clinical or organ-level toxicity at this dose (*48, 49*). Although higher doses (up to 2g/L) have been reported, maintaining hydration and physiological stability is essential in transplantation studies, especially when working with immunocompromised mice.

### *In vivo* vascularization and viability imaging

Functional graft revascularization was assessed as described(*3*) at the indicated days after transplantation by a two-photon microscopy equipped with an incubation system with temperature control (Leica SP5 TPLSM, Leica Microsystems, Wetzlar, Germany). At the time of each experiment, anesthetized mice received an intravenous injection of 80 mg/Kg of rhodamine B isothiocyanate-dextran (RITC-dextran, functional vascular tracer; Sigma) and 2.5 μg/Kg of Hoechst (nuclear tracer; Invitrogen). First, optical fluorescence was used to localize CFDA-positive grafts. Then, stacks of the grafts were acquired every 0,29 µm using a 25x water immersion objective (HCX IR APO L 25x, Numerical Aperture 0,95, Leica), resonant scanner at 8000 lines and blue diode (405 nm), Argon (488 nm) and diode pumped solid state (561 nm). The confocal pinhole was set to its maximum opening. By the end of the visualization, animals were euthanized to perform eye enucleation for posterior immunofluorescence analysis. Collected images were analyzed using ImageJ software (v1.49b; Wayne Rasband, National Institutes of Health, USA). Vascularization was quantified using two parameters. Vascularization area (%) was calculated as the percentage of the graft area occupied by blood vessels, defined as the total area positive for RITC-dextran relative to the total CFDA+ graft area, which reflects the overall extent of vascular coverage within the graft (integrating vessel number and calibre). Vessel density was determined as the number of discrete blood vessels per unit graft area and reflects the abundance of individual vascular structures within the graft, consistent with increased formation of new capillary branches rather than vessel dilation or maturation.

### Gene expression analysis

For gene expression assays, total RNA was isolated using NucleoSpin^®^RNA (Macherey-Nagel Düren, Germany) following the manufacturer’s manual. First-strand cDNA was prepared using Superscript III Reverse Transcriptase (Invitrogen) or High-Capacity cDNA Reverse Transcription Kit (Applied Biosystems, Thermo Fisher Scientific). Real time PCR was performed on an ABI Prism 7900 or Applied Biosystems QuantStudio™ 5 Real-Time PCR system using Gotaq master mix (Promega, Madison, WI, USA). Expression relative to the housekeeping gene *TBP* was calculated using the delta(d)Ct method and expressed as 2^(-dCT). Primer sequences are provided in Table S2.

### Assessment of insulin secretion

To evaluate glucose-stimulated insulin secretion *in vitro*, iBeta-like or iBeta-like/HUVEC spheroids were first washed with phosphate-buffered saline (PBS) and pre-incubated in Krebs-Ringer Buffer containing 2 mM glucose for 45 minutes at 37^°^C. Following this equilibration step, spheroids were sequentially incubated in Krebs Buffer with low glucose (2 mM) and subsequently with high glucose (20 mM) for 60 minutes at 37^°^C. Supernatants were collected after each incubation period for insulin quantification.

For *in vivo* assessment of glucose-induced insulin secretion, mice were fasted for 6 hours prior to receiving an intraperitoneal glucose injection (3g / Kg of body weight). Blood samples were collected a time 0- and 20-min post-injection for insulin measurement.

Human insulin concentration in both *in vitro* supernatants and mouse plasma was determined using a Human Insulin ELISA kit (Crystal Chem, Houston, TX, USA), following the manufacturer’s instructions.

### Immunofluorescence

The eyes containing grafts were fixed overnight in 2% paraformaldehyde and subsequently embedded in paraffin. Each graft was fully sectioned, and the sections were distributed across two sets of 10 slides. Each slide contained 5 columns, with 3 sections per column, spaced 3 μm apart. For immunofluorescence analysis on paraffin sections, a minimum of 10 sections per graft (spanning 2 slides), spaced 90 μm apart, were selected. Tissue sections were rehydrated and subject to heat-mediated antigen retrieval in citrate buffer, permeabilized using 1% Triton X-100 and blocked in 5% normal donkey serum. Then, grafts were incubated at 4^°^C overnight with primary antibodies and for 2 h at room temperature with fluorescent-labelled secondary antibodies (Table S3). For the apoptosis analysis, DeadEnd Fluorometric TUNEL System (Promega) was performed according to the manufacturer instructions. In all cases, images were acquired in a Leica TCS SPE confocal microscope, using a 40x oil immersion objective. Images of positive cells and areas were quantified by using ImageJ software.

### *In toto* staining

After collection from the plate, islets/spheroids were transferred into a 1.5 mL Eppendorf tube, washed, and fixed in 4% paraformaldehyde for 15 minutes at 4 °C. Immunofluorescence staining of whole islets/spheroids was performed in suspension under a stereomicroscope. Samples were permeabilized with 0.8% Triton X-100 and blocked using a solution containing 10% (v/v) FBS and 5% (v/v) normal donkey serum. For subsequent washes, 0.3% (v/v) Triton X-100 was used, and antibody solutions (Table S3) were prepared with 5–10% (v/v) FBS. Following staining with indicated antibodies, samples were mounted on a chamber slide to preserve their three-dimensional structure and imaged using a Leica TCS SPE confocal microscope. Z-stack images were acquired, and sections were analyzed at 5 μm intervals using ImageJ software.

### Cell viability

To assess iBeta-like cell viability, cell spheroids were labeled with 10 µM CFDA SE and 2.5 µg/mL propidium iodide (PI; Invitrogen) for 15 minutes at 37^°^C. Spheroids were then visualized using Leica TCS SPE confocal, and images were analyzed using ImageJ software. Cell viability was calculated as the percentage of CFDA^⁺^PI^⁻^ cells relative to the total number of cells within each spheroid.

HUVEC viability was assayed by MTT assay. In brief, HUVECs were seeded in 96-well plates (10,000 cells/well). After 24 hours, the medium was replaced with fresh EGM-2 containing VEGFA (2 ng/mL), NaW (0.2 mM), both, or neither (control). After 48 hours of treatment, cells were incubated with 0.75 mg/mL MTT (3-(4,5-dimethylthiazol-2-yl)-2,5-diphenyltetrazolium bromide) for 3 hours at 37^°^C. Formazan crystals were dissolved in isopropanol with 0.04 N HCl, and absorbance was measured at 575 and 650 nm using a TECAN Infinite^®^ 200 PRO reader. Corrected OD (OD575–OD650) was normalized to the control (100%).

### Evaluation of angiogenic properties in HUVECs

#### Proliferation assay

HUVECs were seeded at 10,000 cells/well in 96-well plates. After 24 h, the medium was replaced with EGM-2 supplemented with VEGFA (2 ng/mL), NaW (0.2mM), both, or neither (control). Cell proliferation was evaluated at 36 and 72 h post-treatment. At each time point, cells were trypsinized, neutralized with DMEM containing 10% fetal bovine serum (FBS), and counted using the Countess™ Automated Cell Counter (Invitrogen).

#### Migration assay

Cell migration was assessed following established wound healing protocols(*50*). HUVECs were seeded at 250,000 cells/well in 12-well plates and cultured to 90–100% confluence. A scratch was made using a sterile pipette tip, and cells were treated with NaW (0.2mM), VEGFA (2 ng/mL), both, or control. Migration into the wound area was monitored at 0, 2, 4, 6, 8, and 24 h using an inverted optical microscope (OLYMPUS IX51, Hamburg, Germany). Images were analyzed with ImageJ software to quantify the percentage of wound closure relative to the initial scratch area.

#### Tube formation assay

Endothelial tube formation was assessed following established protocols (*51, 52*). A total of 150 µL/well of reduced Growth Factor Basement Membrane Extract (Biotechne, Minneapolis, MN, USA) was added to 48-well plates and incubated at 37^°^C with 5% CO_₂_ for 30 min to allow gelation. HUVECs (40,000 cells/well) were then seeded in conditioned medium containing NaW (0.2mM), VEGFA (2 ng/mL), both, or control. After 3–5 h of incubation, tube-like structures were imaged using an inverted optical microscope (OLYMPUS IX51, Hamburg, Germany). Five representative images per well (center and cardinal axes) were analyzed using the ImageJ Angiogenesis Analyzer macro(*53*) to quantify tube formation parameters.

### Immunoblot analysis

250,000 HUVEC cells/well were seeded in 6-well plates and cultured overnight (15 h) in low serum DMEM (0.2% FBS). Cells were treated for 10 min with NaW, VEGFA (25 ng/mL), or both. Cells were lysed in triple detergent buffer (50 mM Tris-HCl, 150 mM NaCl, 0.1% SDS, 1% NP40, 0.5% sodium deoxycholate) with phosphatase and protease inhibitors (Roche). Lysates were frozen/thawed twice and sonicated (5 × 30 s bursts at 20 kHz). Debris was pelleted and protein quantified using the Lowry assay (Bio-Rad). 15 μg of protein per replicate was separated on 4–12% gradient gels (Bio-Rad) and transferred to PVDF membranes (Perkin Elmer). Membranes were incubated overnight at 4°C with antibodies against ERK1/2 (1:1000, 42/44 kDa), phospho-ERK1/2 (1:1000, 42 kDa), and α-TUBULIN (1:1000, 52 kDa; loading control). Detection was performed using HRP-conjugated secondary antibodies (1:5000, GE Healthcare) and ECL reagent (Pierce). Bands were visualized with LAS4000 (Fuji) and quantified using ImageJ. Antibody information is provided in Table S3.

### Statistical analysis

Data were analysed using GraphPad Prism version 8.00 for Windows (www.graphpad.com) and expressed as the mean ± standard error of the mean (SEM). Student’s t-tests were used for comparison of two groups. One-way ANOVA or Two-way ANOVA were used according to the appropriate number of comparisons followed by recommended post-hoc tests (Tukey’s or Dunnett’s), applied to determine significant differences among multiple comparisons. Differences were considered significant at P < 0.05. P values greater than 0.05 and lower than 0.1 were considered trends.

## Acknowledgements

We gratefully acknowledge Shahrzad Chitgaran and Candice Fournier for their valuable technical support in conducting experiments with HUVEC cells. We thank Solja Eurola, Hossam Montaser and Suvi Tikkakoski for their expert technical support in conducting SC-islet related experiments. We also thank the Advanced Optical Microscopy Unit of the Universitat de Barcelona (CCiTUB) for their assistance with *in vivo* two-photon imaging.

