## Supplemental data for "Sodium tungstate promotes vascularization to support beta cell replacement in diabetes"

Contains:

Figures S1-28
Tables S1-S3

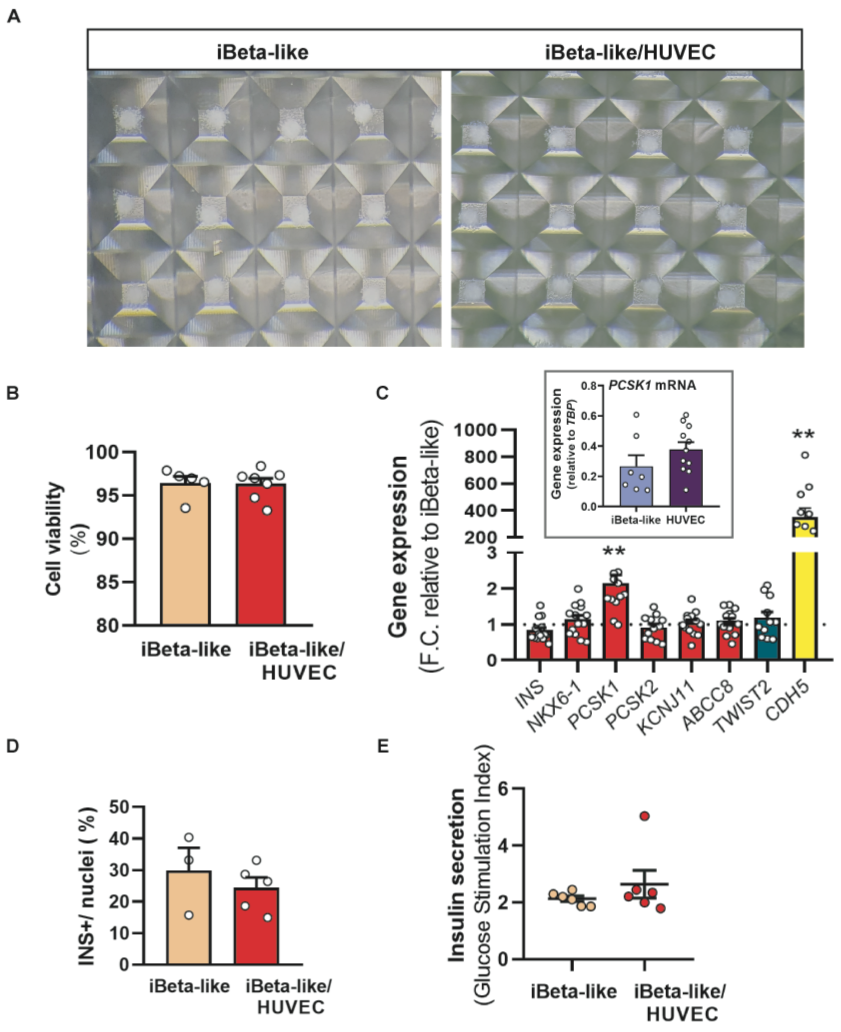


**Supplementary figure 1. Characterization of iBeta-like and iBeta-like/HUVEC spheroids.**

**(A)** Photographs of AggreWell plates showing spheroids composed of iBeta-like or iBeta-like/HUVEC cells five days after seeding single cells into microwells.
**(B)** Cell viability as assessed by CFDA and propidium iodide (PI) staining. Cell viability was calculated as the percentage of $CFDA^{+}PI^{-}$ cells relative to the total number of cells. Each dot represents an individual spheroid.
**(C)** Gene expression analysis by qPCR for the indicated genes in iBeta-like/HUVEC spheroids. Data are expressed as fold-change relative to iBeta-like spheroids, which were set to 1. Dots correspond to eight to ten independent reprogramming experiments. Inset shows *PCSK1* gene expression measured by qPCR in iBeta-like cells and HUVECs. Expression values are normalized to the housekeeping gene *TBP*. Each dot represents an independent spheroid preparation or HUVEC culture
**(D)** Percentage of INS+ cells in the indicated spheroid types at the end of the reprogramming protocol. For iBeta-like/HUVEC spheroids, CDH5+ cells (corresponding to HUVEC cells) were substrated from total nuclei count. Eeach dot represents a different reprogramming experiment.
**(E)** Insulin secretion measured in static incubation assays. The Glucose Stimulation Index represents the fold-change in insulin secretion between high (20 mM) and low (2 mM) glucose conditions. Dots are different spheroids pools from at least three independent reprogramming experiments.
Data are presented as mean ± SEM for the number of n indicated by dots. Statistical analysis was performed using Student's t-test in B,D,E; no significant differences were observed.
In C, ** $p < 0.005$ compared to iBeta-like cells, as determined by one-sample t-test.

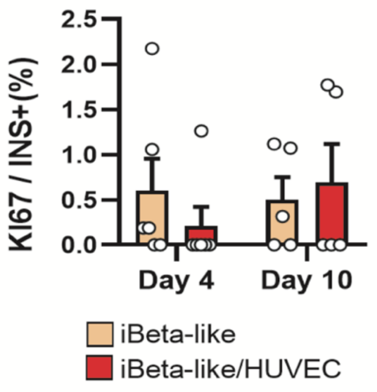


**Supplementary figure 2. Proliferation of iBeta-like cells after transplantation.**

Cell proliferation was assessed as the percentage of $INS^+$ cells co-expressing Ki67, relative to the total number of $INS^+$ cells, in grafts of the indicated types. Data are presented as mean ± SEM for the number of n indicated by dots, with each dot representing an individual graft. Statistical analysis was performed using two-way ANOVA; no significant differences were observed.

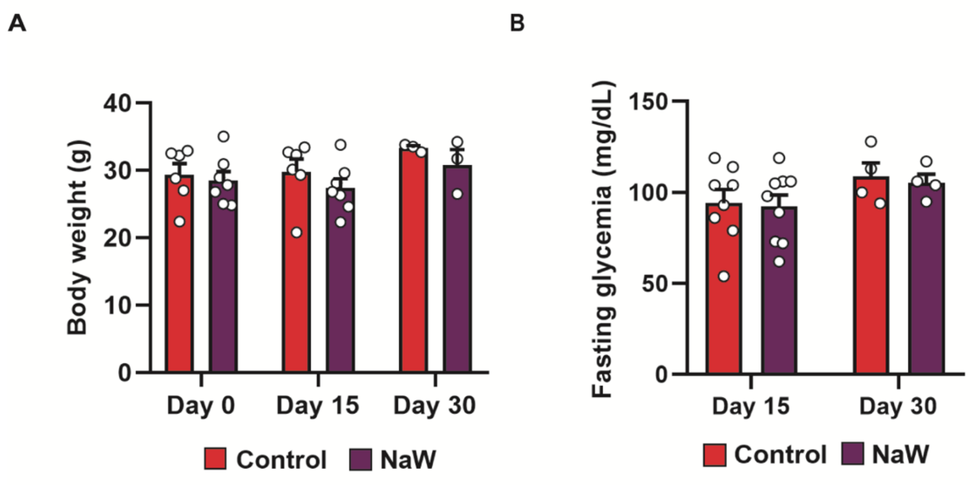


**Supplementary figure 3. Effects of NaW treatment on body weight and glycemia in mice transplanted with iBeta-like/HUVEC spheroids.**

**(A)** Body weight at the day of transplantation (day 0) and on days 15 and 30 post-transplantation in control and NaW-treated mice

**(B)** Fasting glycemia at days 15 and 30 post-transplantation in control and NaW-treated mice.

Data are expressed as mean ± SEM, with each dot representing an individual animal. Statistical analysis was performed using two-way ANOVA, which revealed no significant differences.

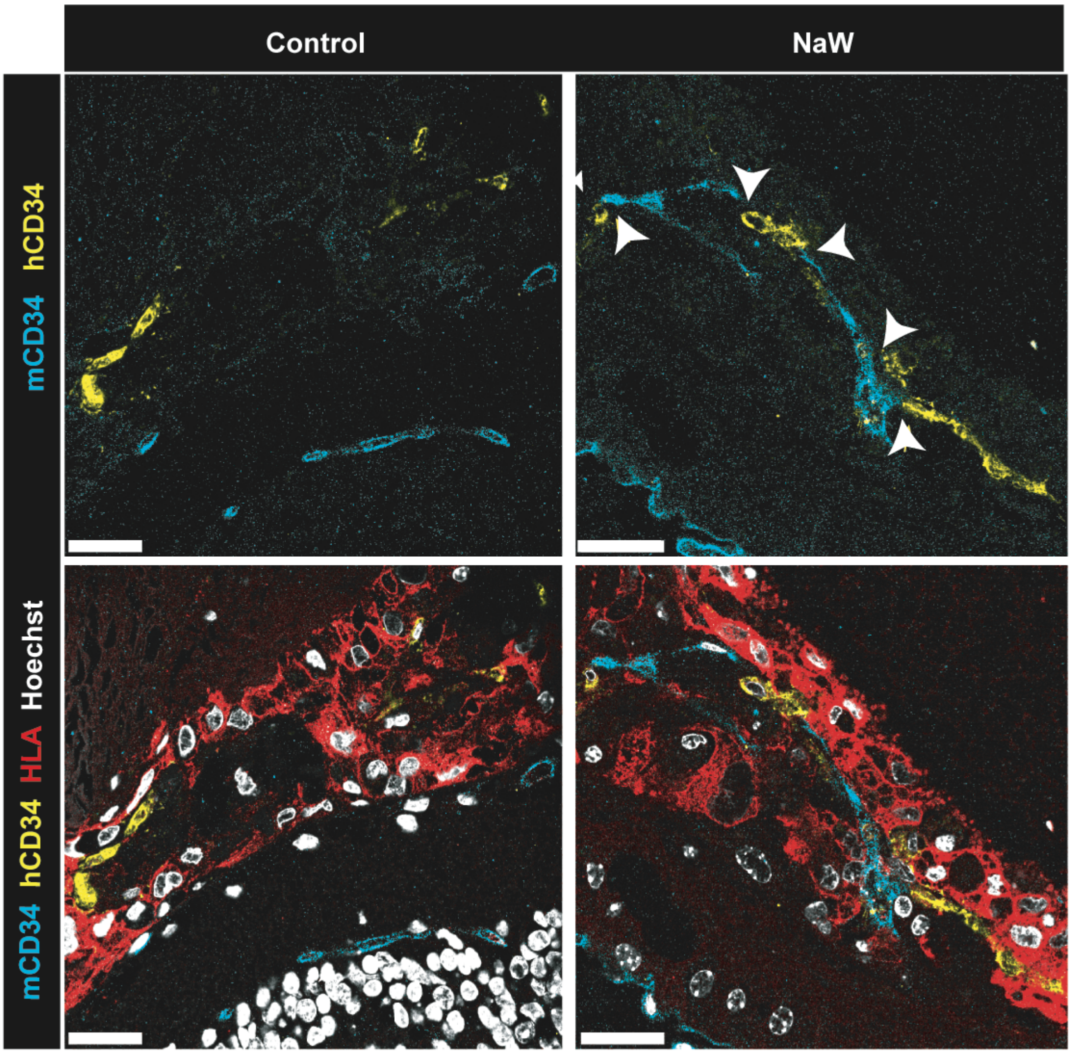


**Supplementary figure 4. Histological characterization of HUVEC and mouse ECs in iBeta-like/HUVEC grafts at day 10 post-transplantation.**

Representative confocal images of iBeta-like/HUVEC grafts at day 10 post-transplantation in control and NaW-treated mice. Grafts were stained for mouse CD34 (cyan; mouse recipient endothelial cells), human CD34 (yellow; HUVECs), human HLA (red; human cells), and Hoechst (blue; cell nuclei). White arrowheads indicate regions where mouse endothelial cells are adjacent to human endothelial cells. Scale bar: 25 μm.

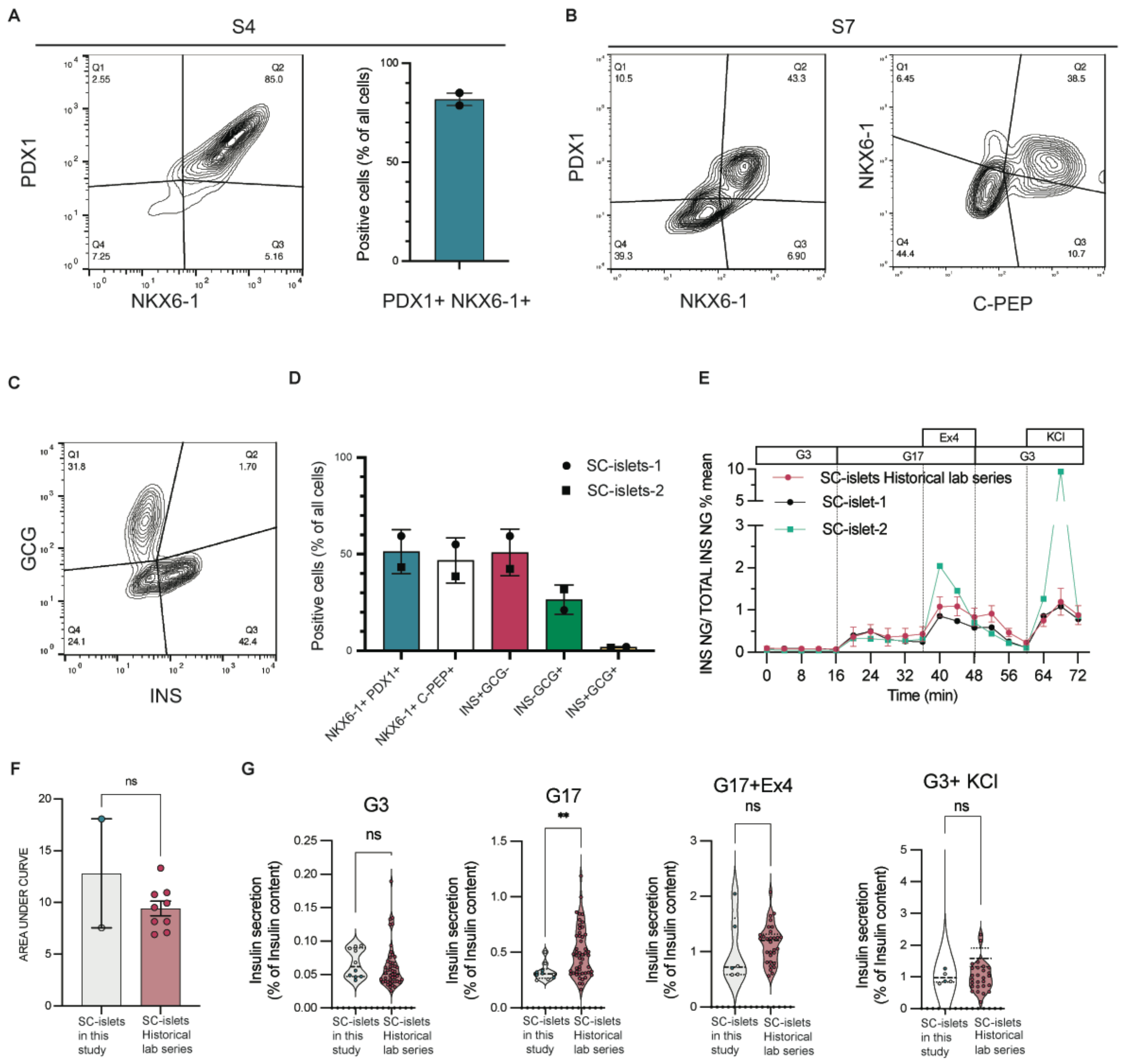


**Supplementary figure 5. Basic characterization of SC-islet differentiation.**

**(A)** Flow cytometry analysis with PDX1 and NKX6-1 antibodies in pancreatic progenitor stage (S4).
**(B,C)** Flow cytometry analysis of SC-islets at S7 week 3 with PDX1 + NKX6-1, NKX6-1+ C-PEP antibodies (B) and INS + GCG antibodies (C).
**(D)** Quantification of the populations from two H1 differentiations (SC-islets-1, SC-islet-2) used in this study.
**(E)** Functional analysis done as dynamic glucose stimulated insulin secretion assay in low glucose (G3), high glucose (G17), high glucose +Exendin4 (Ex4) and low glucose +KCl presented as % of protein content.
**(F)** Area under curve quantifications of E
**(G)** Quantifications of each experimental condition using data points. Comparison of the SC-islet preparations used in this study with laboratory's historical series.

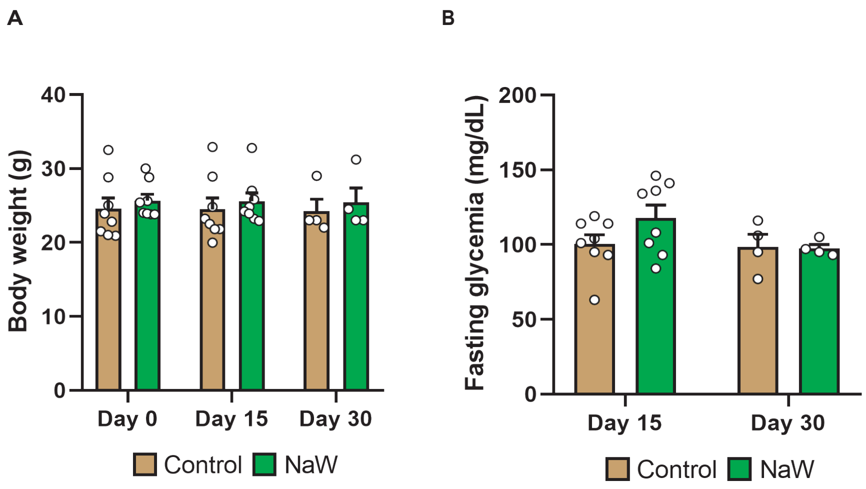


**Supplementary figure 6. Effects of NaW treatment on body weight and glycemia in mice transplanted with SC-islets.**

**(A)** Body weight at the day of transplantation (day 0) and on days 15 and 30 post-transplantation in control and NaW-treated mice
**(B)** Fasting glycemia at days 15 and 30 post-transplantation in control and NaW-treated mice.
Data are expressed as mean ± SEM, with each dot representing an individual animal. Statistical analysis was performed using two-way ANOVA, which revealed no significant differences.

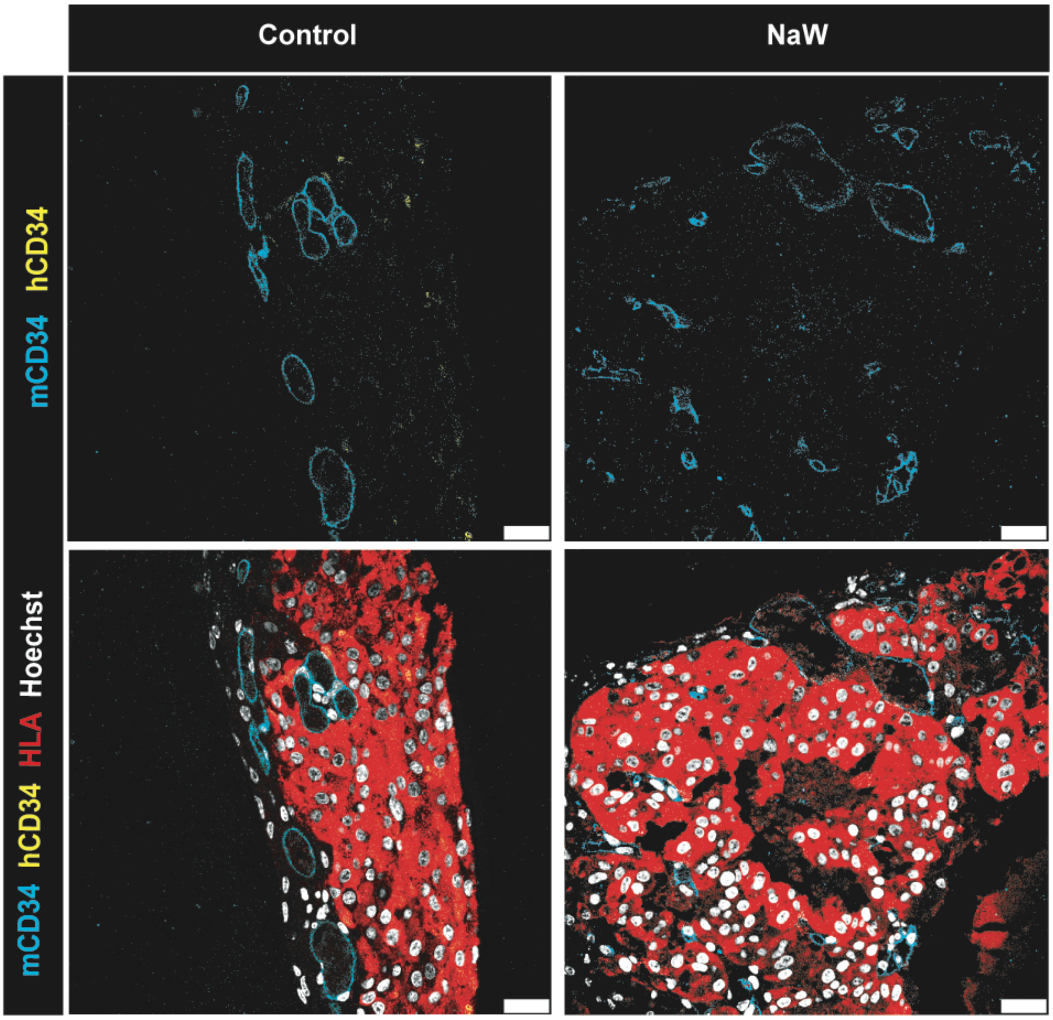


**Supplementary figure 7. Histological characterization of human and mouse ECs in SC-islet grafts at day 15 post-transplantation.**

Representative confocal images of SC-islet grafts at day 15 post-transplantation in control and NaW-treated mice. Grafts were stained for mouse CD34 (cyan; mouse recipient endothelial cells), human CD34 (yellow; HUVECs), human HLA (red; human cells), and Hoechst (blue; cell nuclei). Human $CD34^{+}$ cells were not detected in any samples from either treatment group. Scale bar: 25 μm

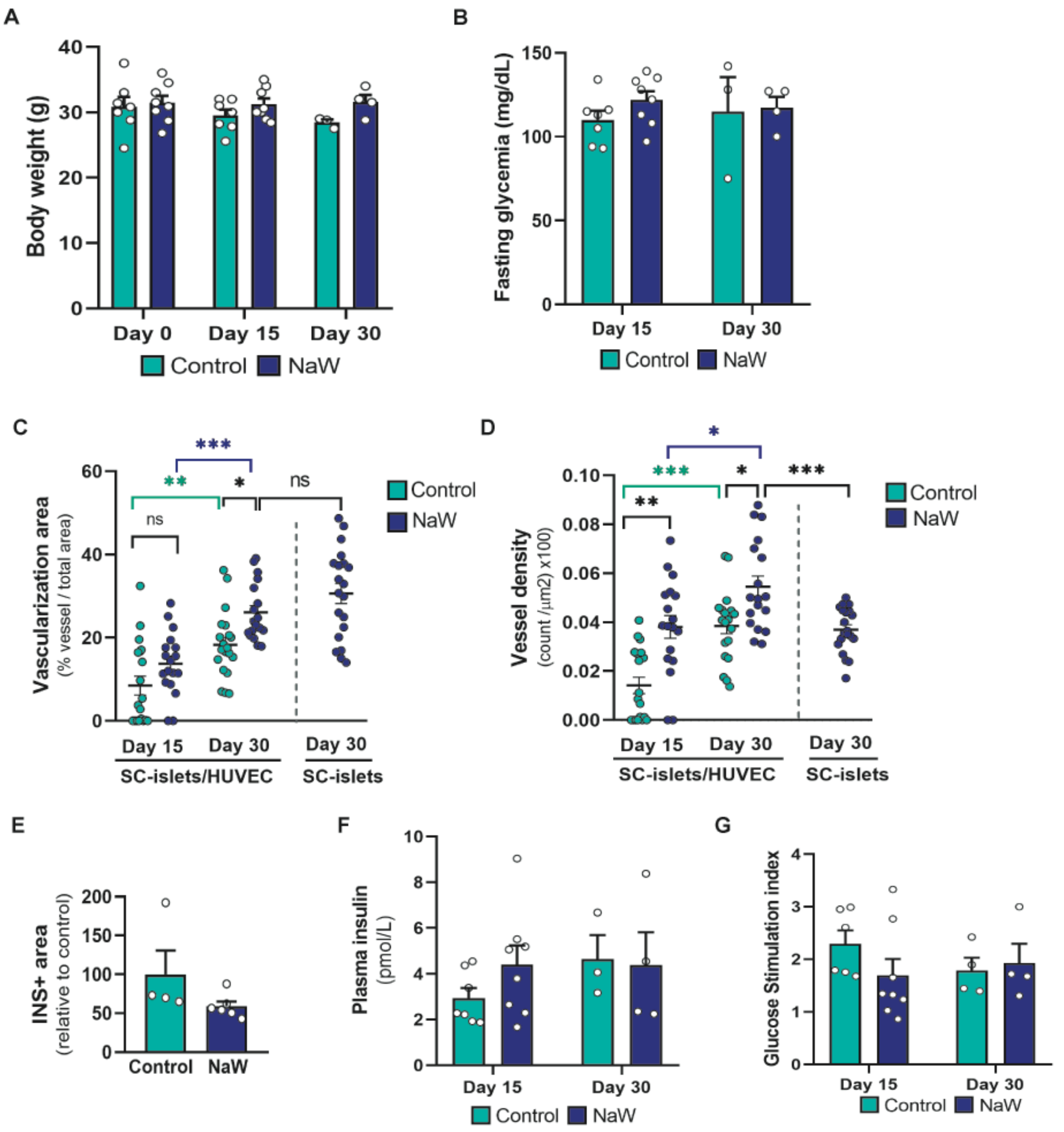


**Supplementary figure 8. Effects of NaW treatment on SC-islet/HUVEC grafts.**

**(A)** Body weight at the day of transplantation (day 0) and on days 15 and 30 post-transplantation in control and NaW-treated mice

**(B)** Fasting glycemia at days 15 and 30 post-transplantation in control and NaW-treated mice.

**(C,D)** Quantitative analysis of vascularized area (C) and vessel density (D) in grafts at days 15 and 30 post-transplantation. For comparative purposes, data obtained from SC-islet grafts in NaW-treated mice at day 30 (Figure 5B,C) are presented.

**(E)** Quantification of insulin-positive area in grafts from control and NaW-treated mice at day 30 post-transplantation. Values are expressed relative to control grafts, with values set to 100%.

**(F)** Fasting human plasma insulin in control mice and mice treated with NaW at days 15 and 30 post-transplantion

**(G)** Fold-increase in plasma insulin 20 min after intraperitoneal glucose administration in control mice and mice treated with NaW at days 15 and 30 post-transplantion.
Data are presented as mean ± SEM for the number of n indicated by dots. In A,B, E, F: dots represent individual mice. In C,D: dots represents one measured area, from a total of n= 5-7 grafts.. In G: dots represent the mean value from individual grafts. Statistical analysis was performed using two-way ANOVA (A-D, F,G) and unpaired Student's t-test (E, and C;D for the comparison SC-islets and SC-islet/HUVEC at day 30): *p < 0.05, **p < 0.005, *** p<0.0001 between the indicated conditions. In C-D, significance bars and symbols are color-coded for comparisons within the same treatment group across different days and shown in black for comparisons between different treatment groups and/or graft types.

**Supplementary Table S1.**

Human islet donor information.

| Unique identifier | HM102 | HM116 |
|---|---|---|
| **Donor age (years)** | 70 | 61 |
| **Donor sex (M/F)** | M | F |
| **Donor BMI (kg/m²)** | 24,7 | 28,5 |
| **Donor HbA1c** | 5.6 | 5,3 |
| **Islet isolation centre** | Montpellier | Montpellier |
| **Donor history of diabetes** | No | No |
| **Diabetes duration (years)** | - | - |
| **Glucose-lowering therapy at time of deathc** | - | - |
| **RECOMMENDED INFORMATION** | | |
| **Donor cause of death** | Stroke | Stroke |
| **Warm ischaemia time (h)** | <2h | <2h |
| **Cold ischaemia time (h)** | 11h45 | 8h40 |
| **Estimated purity (%)** | 85 | 85 |
| **Estimated viability (%)** | >80 | >80 |
| **Total culture time (h)d** | 3 days | > 2 days |
| **Glucose-stimulated insulin secretion or other functional measuremente** | GSIS ratio (16.7 mM vs 2.8mM) = 1,71 | ? |
| **Handpicked to purity? Please select yes/no from drop down list** | ? | ? |
| **Additional notes** | anti-thrombosis / hypertension and anticholesterol treatments | hypertension treatment |

**Supplementary Table S2.**

List of primers used for gene expression analysis

| Gene | | Sequences |
|---|---|---|
| ***TBP*** | Forward<br>Reverse | 5’ - ATCCCTCCCCCATGACTCCCATG -3’<br>5’ - ATGATTACCGCAGCAAACCGC -3’ |
| ***INS*** | Forward<br>Reverse | 5’ - GCAGCCTTTGTGAACCAACA -3’<br>5’ - TTCCCCGCACACTAGGTAGAGA |
| ***NKX6-1*** | Forward<br>Reverse | 5’ - ACACGAGACCCACTTTTTCCG -3’<br>5’ - GCCCCGCCAAGTATTTTGTT -3’ |
| ***PCSK1*** | Forward<br>Reverse | 5’ - AAGCAAACCCAAATCTCACCTGGC -3’<br>5’ - TCACCATCAAGCCTGCTCCATTCT -3’ |
| ***PCSK2*** | Forward<br>Reverse | 5’ - CCGGGTTCCTCTTCTGTGTC -3’<br>5’ - AGCAAAGGGAAGCTTTCGGA-3’ |
| ***KCNJ11*** | Forward<br>Reverse | 5’ - TGTGTCACCAGCATCCACTC -3’<br>5’ - CACTTGGACCTCAATGGAGAA -3’ |
| ***ABCC8*** | Forward<br>Reverse | 5’ - AGACCCTCATGAACCGACAG -3’<br>5’ - GGCTCTGTGGCTTTTCTCTC -3’ |
| ***TWIST2*** | Forward<br>Reverse | 5’ - CGCAAGTGGAATTGGGATGC -3’<br>5’ - CGATGTCACTGCTGTCCCTT -3’ |
| ***CDH5*** | Forward<br>Reverse | 5’ - ATGCGGCTAGGCATAGCATT -3’<br>5’ - TGTGACTCGGAAGAACTGGC -3’ |
| ***VEGFA*** | Forward<br>Reverse | 5’ - AGTCCAACATCACCATGCAG -3’<br>5’ - TTCCCTTTCCTCGAACTGATTT -3’ |

**Supplementary Table S3**

Antibody inventory with corresponding techniques
(IF: immunofluorescence staining in paraffin sections; IT: in toto immunofluoresce staining; WB: immunoblot)

| Primary antibody | Target | Raised in | Working dilution | Commercial source | Technique |
|---|---|---|---|---|---|
| **Anti-Insulin** | Human, mouse | Guinea Pig | 1/500 | Dako | IF |
| **Anti-Glucagon** | Human, mouse | Mouse | 1/250 | Invitrogen | IF |
| **Anti-HLA Class 1 ABC** | Human | Mouse | 1/100 | Abcam | IF |
| **Anti-C-Peptide** | Human | Rat | 1/300 | Hybridoma Bank | IF, IT |
| **Anti-Ki67** | Human, mouse | Rabbit | 1/200 | Invitrogen | IF |
| **Anti-CD34** | Mouse | Rat | 1/150 | eBioscience | IF |
| **Anti-CD34** | Human | Sheep | 1/20 | Biotechne | IF |
| **Anti-veCADHERIN** | Human, mouse | Rabbit | 1/150 | Abcam | IF, IT |
| **Anti-VEGF** | Human, mouse | Rabbit | 1/200 | Millipore | IF |
| **Anti-P-p44/42 MAPK (T202/Y204)** | Human, mouse | Rabbit | 1/1000 | Cell Signaling | WB |
| **Anti-p44/42 MAP Kinase** | Human, mouse | Rabbit | 1/1000 | Cell Signaling | WB |
| **Anti-alpha Tubulin** | Human, mouse | Mouse | 1/1000 | Sigma | WB |

| Secondary antibody | Target | Raised in | Working dilution | Commercial Source | Technique |
|---|---|---|---|---|---|
| **AlexaFluor 488** | Rabbit | Donkey | 1/250 | Jackson Immunoresearch, | IF |
| **AlexaFluor 488** | Guinea Pig | Donkey | 1/250 | Jackson Immunoresearch, | IF |
| **Cy2** | Rat | Donkey | 1/250 | Jackson Immunoresearch, | IF |
| **Cy2** | Sheep | Donkey | 1/250 | Jackson Immunoresearch, | IF |
| **AlexaFluor 555** | Mouse | Donkey | 1/400 | Molecμlar Probes | IF |
| **AlexaFluor 555** | Rabbit | Donkey | 1/400 | Molecμlar Probes | IF, IT |
| **AlexaFluor 647** | Rat | Donkey | 1/250 | Jackson Immunoresearch | IF, IT |
| **AlexaFluor 647** | Rabbit | Donkey | 1/250 | Jackson Immunoresearch | IF |
| **Anti-IgG Peroxidase** | Mouse | Sheep | 1/5000 | GE Healthcare | WB |
| **Anti-IgG Peroxidase** | Rabbit | Donkey | 1/5000 | GE Healthcare | WB |
